# Cell cycle progression defects and impaired DNA damage signaling drive enlarged cells into senescence

**DOI:** 10.1101/2022.09.08.506740

**Authors:** Sandhya Manohar, Marianna E. Estrada, Federico Uliana, Gabriel E. Neurohr

## Abstract

Cellular senescence plays an important role in development, ageing, and cancer biology. Senescence is associated with increased cell size, but how this contributes to permanent cell cycle exit is poorly understood. Using reversible G1 cell cycle arrests combined with growth rate modulation, we examined the effects of excess cell size on cell cycle progression in human cells. We show that enlarged cells paradoxically have high levels of G1/S regulators relative to cells that were maintained at physiological size but also induce p21, which restrains cell cycle entry and protects against cell division failure. Furthermore, we find that enlarged cells bear an increased propensity for DNA breakage and concomitant DNA damage repair defects that are established during G1. Based on these observations, we propose that impaired DNA damage repair pathways prime enlarged cells for persistent replication-acquired damage, ultimately leading to catastrophic cell cycle failure and permanent cell cycle exit.

## Introduction

Cellular senescence describes a permanent state of cell cycle arrest induced by exogenous or endogenous stressors [1, 2]. These include but are not limited to telomere attrition, genotoxic and proteotoxic stress, and oncogene activation. In addition, many cancer therapies (e.g., chemotherapies and radiotherapies) trigger senescence and thus permanent cell cycle withdrawal in tumor cells [3]. Still, though senescence suppresses proliferation, tumors containing persistent senescent cells can be more invasive and are associated with worse outcomes [3, 4]. Thus, identifying how short-term cellular insults can cause a durable loss of proliferative potential is essential for understanding the benefits and limitations of senescence induction as an antitumor therapeutic strategy.

Despite being caused by diverse stimuli, it has long been observed that senescent cells are larger than cycling cells [5]. This observation is important because—although healthy mammalian cells exist at a wide range of sizes—the cell size distribution for a given cell type is typically narrow [6]. Deviations from this range are associated with a loss of fitness, cell cycle failure, and permanent cell cycle arrest [7, 8]. More recent work has demonstrated that increased cell size is sufficient to withdraw cells from the cell cycle [7-12]. Still, it is unclear what pathways drive cell cycle exit in excessively large cells and how increased size activates these pathways. Answering these questions will provide key insight into how permanent cell cycle arrest is conferred in senescent cells.

In mammalian cells, senescence induction typically requires one of two signaling axes: the Rb pathway or the p53/p21 pathway [13]. Rb is a cell cycle inhibitor that binds E2F-family transcription factors to inhibit the transcription of G_1_/S cyclins and components of the DNA replication apparatus [14], thereby blocking cell cycle entry. In the classical model, Rb is partially inactivated by Cdk4/6-mediated phosphorylation during late G_1_. This facilitates low levels of E2F-mediated transcription, a target of which is cyclin E. Cyclin E production activates Cdk2 (the G_1_/S cyclin-dependent kinase) which further phosphorylates Rb to release E2F transcriptional inhibition completely [15]. Rb expression levels also play a role in dictating its activity: high levels of Rb are sufficient to cause G_1_ cell cycle arrest [16] and loss of Rb drives cell cycle progression forward [17, 18]. Furthermore, others have shown that reducing Rb concentrations during growth is a mechanism for linking cell growth to cell cycle entry [19, 20]. Thus, Rb abundance and its phosphorylation state are both important for regulating the G_1_/S transition. The p53/p21 pathway is another major cell cycle arrest pathway in mammalian cells.

p53 is a transcription factor that is activated upon DNA damage [21], aneuploidy [22-24], oxidative damage [25], and other stressors. Following DNA damage, p53 stabilization is mediated by ATM and ATR protein kinases [21, 26, 27]. In this context, many of p53’s transcriptional targets are implicated in DNA damage repair and cell cycle arrest [28]. p21—one of p53’s main transcriptional targets—is a Cdk1/2/4/6 inhibitor that halts cell cycle progression through the G_1_/S boundary [29]. Depending on context, p21 expression can contribute to temporary cell cycle arrest, permanent cell cycle arrest (senescence), apoptosis, or DNA repair [29]. Still, p21 dynamics are complex, and p21 expression can have opposing effects under different circumstances. Indeed, high levels of p21 cause cell cycle arrest, whereas intermediate levels can drive cell cycle progression forward [30, 31]. In addition to directly binding Cdks, p21 can interact with components of the DNA replication apparatus to halt DNA synthesis and modulate repair pathways [32]. p21 transcription can also be stimulated independent of p53, including through the HRAS-Raf-MAPK pathway and by various transcription factors (e.g., SP1, AP2, C/EBPα/β) [29].

There is considerable crosstalk between the p53/p21 and Rb pathways. Because Rb mediated inhibition of E2F is dictated by Cdks, high levels of p21 block Cdk activity and therefore prevent E2F activation. Moreover, Rb directly regulates p53 stability through its interaction with MDM2, a p53-directed ubiquitin ligase [33]. Still, it is yet unclear if increased cell size activates either or both pathways to drive permanent cell cycle exit.

Here, we show that continued cell growth is required to induce long-term cell cycle exit in human cells treated with the Cdk4/6 inhibitor palbociclib. We find that enlarged cells upregulate p21, which protects against cell cycle entry and subsequent cell cycle failure. Indeed, large cells that enter the cell cycle have S/G_2_/M delays and undergo mitotic catastrophes with high frequency. These cell cycle abnormalities are accompanied by a blunted p53 response and DNA damage repair defects in enlarged cells, resulting in a high propensity for DNA damage during G_1_. We propose that these defects prime cells for high levels of replication-acquired damage during S-phase, leading to catastrophic mitoses followed by permanent cell cycle withdrawal. Together, these results provide a framework for defining the fate of enlarged G_1_ cells and show that excess cell size renders cells prone to DNA damage by interfering with DNA damage signaling and repair.

## Results

### Continued cell growth induces permanent cell cycle exit following a prolonged G_1_ arrest

In order to understand how increased cell size influences cell cycle progression, we used the Cdk4/6 inhibitor palbociclib to arrest hTERT-RPE1 (hereafter referred to as RPE1) cells in G_1_ (**Figure S1A**-**S1B**). Because G_1_-arrested cells continue to accumulate biomass [7], this treatment enables continued growth in the absence of division and significantly increases cell size. To disentangle the effects of increased size from those caused by prolonged Cdk4/6 inhibition, we employed two control strategies: (1) seeding cells at high confluence (contact inhibition) prior to palbociclib treatment and (2) inhibiting mTOR activity during G_1_ arrest using the small molecule Torin1 (**Figure 1A**). Using these approaches, we were able to obtain enlarged G_1_-arrested cells and corresponding control cells that were close in size to untreated cells despite experiencing G_1_ cell cycle arrest for the same duration (**Figure 1B, Figure S1C**). Control cells for which growth was restricted during a 6-day palbociclib-mediated arrest using either of these strategies are hereafter called “size-constrained.” Importantly, though cells that were co-treated with Torin1 to constrain cell size could simply be switched to drug-free media to examine cell cycle re-entry, cells that were contact inhibited had to be re-seeded at a lower density to facilitate cell cycle re-entry (**Figure 1A**). In both cases, size-constrained cells were plated in palbociclib alone for one day following arrest to allow them to recover from the effects of Torin1 treatment or contact inhibition respectively. Maintenance of G_1_ arrest was confirmed using FUCCI cell cycle reporters, which show that mAG-geminin^1-110^ (an S/G_2_/M marker) levels are low in 6-day arrested cells relative to cycling cells and are consistent between enlarged and size-constrained cells (**Figure S1A**-**S1B**).

**Figure 1:**
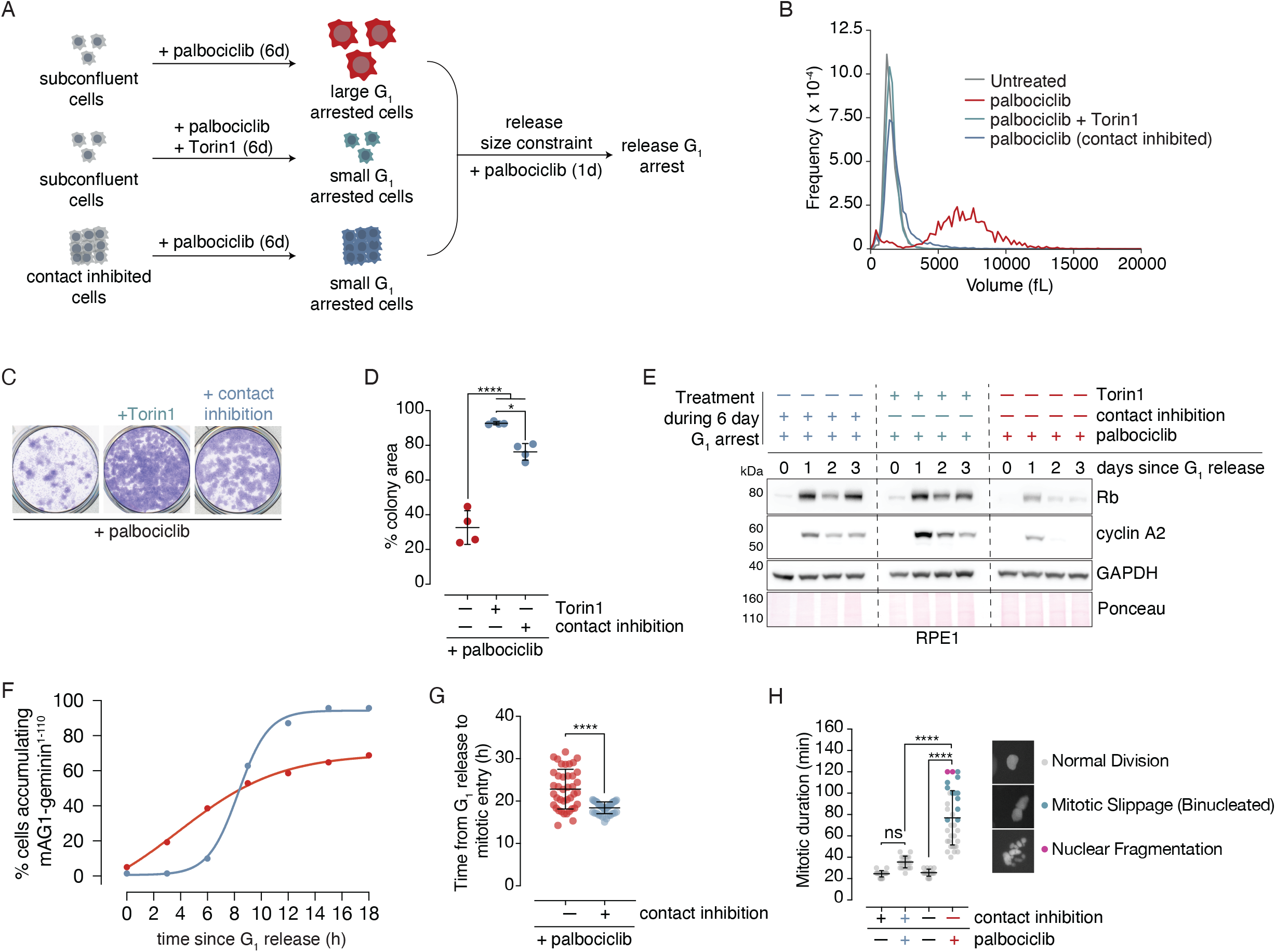
Continued cell growth induces permanent cell cycle exit following a prolonged G_1_ arrest. **(A)** Schematic for constraining cell size in the presence of palbociclib using mTOR inhibition by Torin1 or contact inhibition. To grow cells large, cells were plated and maintained at low density while being treated with palbociclib alone for 6 days. To constrain cell size using Torin1, cells were plated at a low density and treated with palbociclib + Torin1 for 6 days. To constrain cell size using contact inhibition, cells were plated at high density and treated with palbociclib for 6 days. In all cases, media and drugs were replaced every 1-3 days. After 6 days, cells were re-seeded at low density (if necessary) and switched to media containing palbociclib alone for 24 hours before performing G_1_ experiments or releasing cells into fresh media without palbociclib for release experiments. For release experiments, cells were washed three times in media prior to release. **(B)** Coulter Counter-based cell volume measurements for untreated (cycling), palbociclib-treated, palbociclib + Torin1 treated, and palbociclib + contact inhibition RPE1 cells after a 6-day treatment. **(C)** Long-term colony formation assay. RPE1 cells were treated as in (**A**) and for 6 days plus 1 day for recovery and were then seeded at 250 cells/cm^2^ in the absence of drugs for 10 days. Cells were then crystal violet stained to visualize colonies. **(D)** Quantification of (**C**), n = 4. * : p = 0.01, **** : p < 0.0001. p-values were calculated by oneway ANOVA followed by Tukey’s multiple comparisons test. **(E)** Cell cycle release time course following release in size-constrained (contact-inhibited and Torin1-treated) and enlarged RPE1 cells. After 6 days + 1 day of recovery, cells were released into fresh media without palbociclib and collected at the indicated time points. Cell lysates were analyzed by western blot for the indicated protein abundances. GAPDH and Ponceau membrane staining were used as loading controls. **(F)** RPE1 FUCCI cells were treated as in (**A**) and imaged following release from G_1_ arrest. Cells were tracked for 18 hours, and cumulative frequency curves were plotted for cells that had started accumulating mAG1-geminin^1-110^ (the S/G_2_/M marker) based on the fraction of cells at each time point that were above an arbitrary mAG1-geminin^1-110^ intensity threshold. At least 45 cells were analyzed for each condition at each time point. **(G)** The timing from G_1_ release (start of imaging) until mitotic entry for the first 40 cells that reach mitosis for each condition in the experiment described in (**F**). Mitotic entry was defined as the frame at which nuclear envelope breakdown (NEB) occurred. p-value was calculated using a two-tailed unpaired t-test. **** : p < 0.0001. **(H)** For cells that reached mitosis following G_1_ release, mitotic duration (the time from NEB to flattening or division) was quantified, and mitotic fates (normal division, slippage, or nuclear fragmentation) were documented. p-values are given for mitotic duration and were calculated by one-way ANOVA followed by Tukey’s multiple comparisons test. **** : p < 0.0001. ns : p = 0.1. Error bars = mean ± SD.

Previous work has shown that cells that have grown beyond their physiological size range fail to proliferate and enter senescence [7, 8, 34]. In agreement with these observations, we found that constraining cell size using contact inhibition or Torin1 treatment is sufficient to rescue long-term proliferation following G_1_ arrest release in RPE1 cells (**Figure 1C-1D**). To understand at what point following G_1_ arrest release large RPE1 cells undergo cell cycle failure, we monitored cell cycle markers by western blot following release in large cells and cells whose size had been constrained using Torin1 and contact inhibition (**Figure 1E**). We found that size-constrained cells recover and maintain cyclin A2 expression (which is observed in S/G_2_/early-M cells [35]) following release from G_1_ arrest, indicating that they continue to cycle. In contrast, cells that grew large recover cyclin A2 expression only transiently. These data suggest that enlarged RPE1 cells enter the cell cycle once following G_1_ release before permanently exiting the cell cycle.

To further understand the events that lead to cell cycle failure in RPE1 cells, we used live-cell imaging to investigate cell cycle re-entry dynamics. Using the experimental scheme shown in **Figure 1A**, we monitored cell cycle progression in large and size-constrained RPE1 cells expressing FUCCI cell cycle reporters [36] following G_1_ arrest release. Though nearly all size-constrained cells began accumulating mAG1-geminin^1-110^ within 18 hours of release, 40% of enlarged cells remain arrested in G_1_ (**Figure 1F**). In large cells that entered mitosis, mitotic entry was delayed relative to cells that were kept small (**Figure 1G**). Large cells also spent significantly longer in mitosis and had a high frequency of abnormal mitotic outcomes, including nuclear fragmentation and mitotic slippage yielding binucleated cells (**Figure 1H, Figure S1D**). We observed similar post-mitotic defects in fixed RPE1 WT cells, as measured by nuclear staining following release (**Figure S1E**). Thus, enlarged cells experience cell cycle delays and have a propensity for prolonged, erroneous mitoses. The latter two observations are consistent with unresolved DNA replication defects prior to mitotic entry [37]. Indeed, work from others has demonstrated that G_2_/M cells that were released from a palbociclib-mediated arrest enter mitosis with replication-acquired DNA damage [11]. Because we observe these defects only in large cells and not in size-constrained cells, we conclude that the cell cycle failures observed following palbociclib treatment are a consequence of increased cell size and not prolonged G_1_ cell cycle arrest.

### DNA replication machinery is not limiting in enlarged G_1_ cells

Because changes in cell size can profoundly remodel the proteome [12] and cause cytoplasmic dilution [7], we hypothesized that changes in the abundance of factors that control faithful cell cycle progression in large cells could be responsible for the cell cycle defects we observed in enlarged RPE1 cells. In order to address this possibility, we used TMT-based quantitative proteomics to measure relative protein abundances in 6-day palbociclib arrested large cells compared to size-constrained cells (**Figure 1A**). To uncover proteins that change in response to G_1_ arrest duration as opposed to size, we also included a 2-day palbociclib-arrest time point in this experiment (**Figure 2A**). Based on this experimental scheme, we identified and quantified 5884 proteins based on at least two peptides (**Figure S2A**-**S2E, Table S1**). Hierarchical clustering analysis showed that Day 2 and Day 6 palbociclib-arrested samples clustered together, whereas Torin1 and contact inhibited samples clustered together (**Figure S2F**). Thus, the samples that exceeded physiological size clustered together, and cells that were maintained close to physiological size clustered together. Together, these observations suggest that this experimental setup was a viable strategy for stratifying size-related proteome changes from cell cycle arrest-related changes in G_1_ cells.

**Figure 2:**
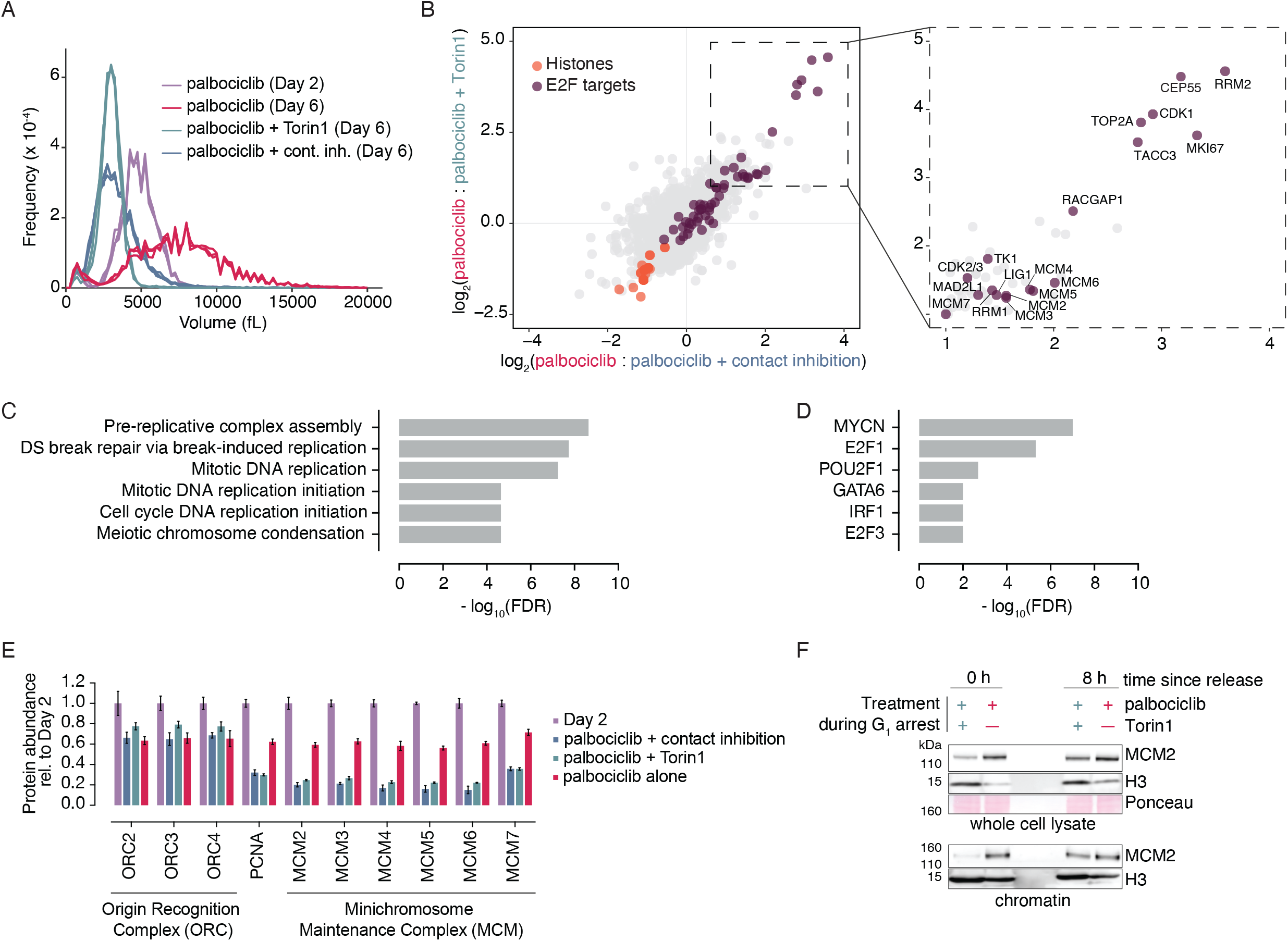
DNA replication machinery is not limiting in enlarged G_1_ cells. **(A)** Coulter Counter-based cell volume measurements for samples (triplicate) used for TMT-based MS experiment. **(B)** (*left*) Comparison of relative protein abundances in enlarged cells vs. either Torin1 (y-axis) or contact inhibited (x-axis) cells plotted as log_2_(fold change). E2F targets are labeled in purple, and histones are labeled in orange. (*right*) Inset showing upregulated proteins in palbociclib-treated cells relative to both size-constrained conditions with a log_2_(fold change) ≥ 1. Proteins encoded by E2F target genes are labeled with their gene names. **(C)** Gene ontology (GO) enrichment of biological processes in the subset of proteins shown in the inset of (**B**). **(D)** Gene ontology (GO) enrichment for transcription factor targets in the subset of proteins shown in the inset of (**B**). **(E)** Protein abundances of origin recognition complex (ORC), PCNA, and minichromosome maintenance complex (MCM) components as measured by mass spectrometry. All measurements are normalized to the mean of the Day 2 time point. Error bars: mean ± SD. **(F)** Western blots of whole cell lysate (*top*) and chromatin fractions (*bottom*) from enlarged and size-constrained cells following an extended G_1_ arrest (0 h) and 8 h after release using antibodies against the indicated proteins. Ponceau staining was used as a loading control for whole cell lysates. H3 is shown to confirm subscaling of histones in whole cell lysate and as a loading control for the chromatin fractions.

In order to identify proteins whose abundances are differentially regulated in large cells relative to both of the size-constrained conditions we used, we compared protein abundances in large cells to each size-constrained condition (**Figure 2B**, *left*). Comparison of both fold changes revealed a linear correlation (adj. R^2^ = 0.4311). We filtered our dataset for proteins that change significantly (adj. *p*-value < 0.05) with the same directionality relative to both size-constrained conditions. Using this strategy, we identified 44 proteins whose abundances decreased by more than 50% in enlarged cells relative to size-constrained cells (**Table S2**). This subset of proteins was comprised mostly of histones (**Figure 2B**), which was expected and served as an internal control given that histone abundance scales with DNA content rather than cell size [12, 38]. In a second step, we filtered our dataset for proteins that increased significantly more than 2-fold in enlarged cells relative to size-constrained cells. This analysis revealed 50 proteins (**Figure 2B**, *inset*, **Table S3**). Gene ontology (GO) analysis for biological processes revealed a strong enrichment in proteins in this subset that are involved in DNA replication and cell cycle progression (**Figure 2C**). Transcription factor regulator relationship analysis revealed that many of these upregulated proteins are known E2F transcriptional targets (**Figure 2B**, *inset*, **Figure 2D**). In addition, many upregulated proteins are positive regulators of the G_1_/S transition (e.g., CDK1/2, CDK4, CCND1, CKS2) (**Figure S2G**). Together, these results suggest that enlarged G_1_ arrested RPE1 cells are not deficient in G_1_/S gene expression relative to size-constrained cells. Of note, our imaging data (**Figure S1A**-**S1B**) show that this is not due to enlarged cells escaping the palbociclib-mediated G_1_ arrest.

Recently published proteomic data indicate that components of the DNA replication apparatus are depleted over time during palbociclib-mediated G_1_ arrest, which could impair origin licensing and cause replication stress and subsequent cell cycle failure upon release [11]. Comparison of our Day 2 and Day 6 palbociclib-arrested cells reproduce this finding, demonstrating a loss in MCM complex components and ORC components as a function of G_1_ arrest duration (**Figure 2E**). Still, we found that components of the MCM complex are *more* abundant in large cells compared to size-constrained cells, though this difference is reduced 8 hours after release from arrest (**Figure 2E-2F**). To address the possibility that enlarged cells are deficient in replisome loading, we measured MCM2 association with chromatin as others have done previously [39]. Mirroring total MCM levels, we found that arrested large cells have more chromatin-associated MCM2 than size-constrained cells, and this difference is eliminated after an 8-hour release (**Figure 2F**, *bottom*). Together, these data suggest that replisome abundance and loading are not limiting for DNA replication in enlarged cells. This finding is consistent with the observation that MCM components are expressed in vast excess of what is required for faithful DNA replication [40].

### Excess G_1_ cell size activates p53-dependent signaling in RPE1 cells

Despite high levels of positive G_1_/S regulators in enlarged G_1_ cells, our proteomic data revealed that levels of the G_1_/S Cdk inhibitor p21 (*CDKN1A*) are elevated in enlarged RPE1 cells, and we confirmed this by western blot (**Figure 3A, Figure S3A)**. In contrast, we found that levels of p16 (*CDKN2A*; another Cdk inhibitor protein) are reduced in enlarged RPE1 cells relative to size-constrained cells (**Figure S3B**-**S3C**). Because p16 negatively regulates a similar set of genes as Rb [41], its depletion could also contribute to the elevated levels of G_1_/S regulators we observed in enlarged cells.

**Figure 3:**
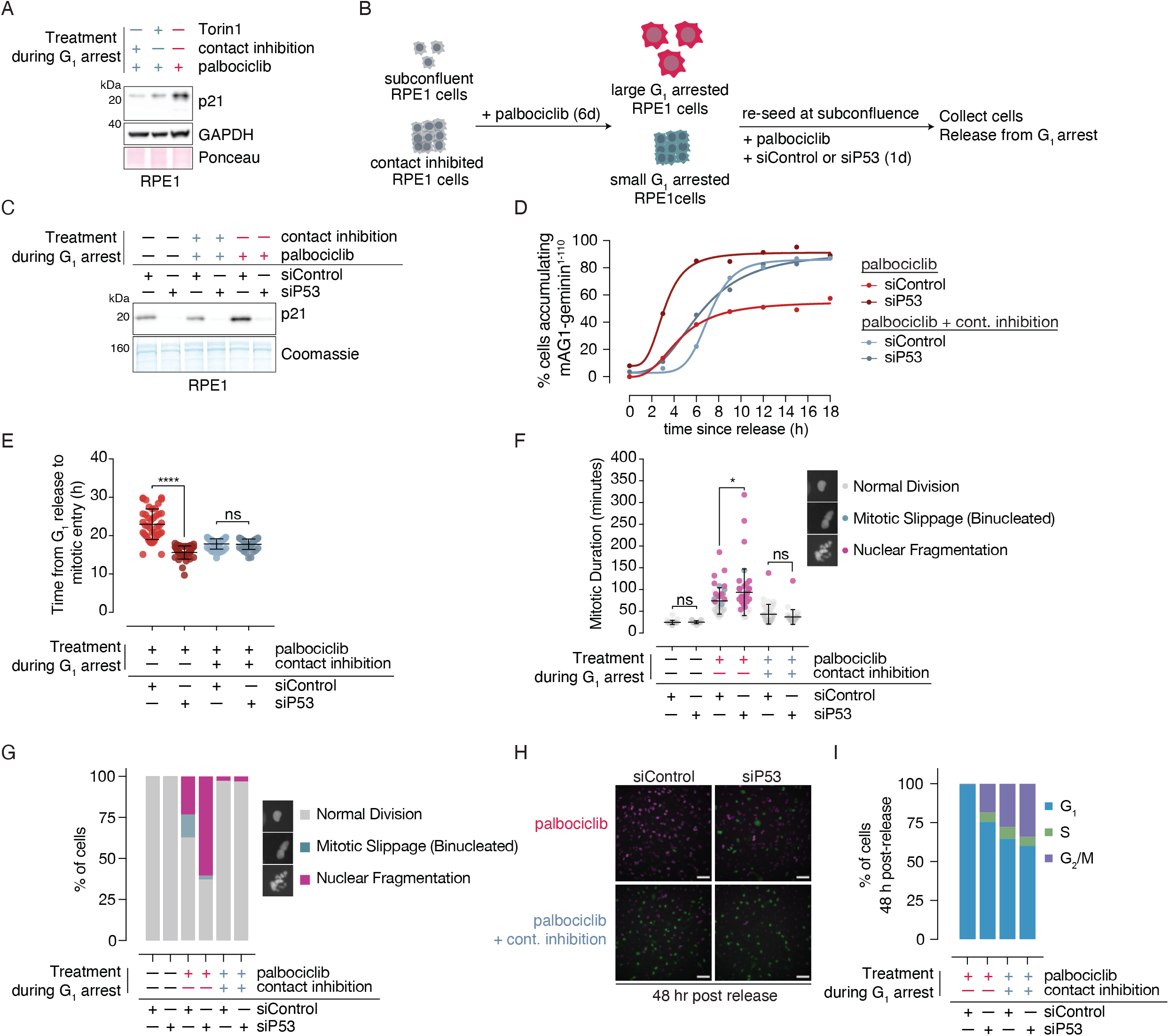
Excess G_1_ cell size activates p53-dependent signaling in RPE1 cells. **(A)** Western blot showing p21 levels in size constrained (Torin1 and contact inhibited) and enlarged G_1_ RPE1 cells. GAPDH and Ponceau staining were used as loading controls. **(B)** Schematic for performing p53 knockdown experiments in enlarged and size-constrained cells. **(C)** Western blot depicting p21 levels in cycling, size-constrained, and enlarged cells with and without p53 knockdown. Note that p53 was not detected by western blot in cycling or G_1_ arrested RPE1 cells. Coomassie staining of the gel was used as a loading control. **(D)** Cumulative frequency curves for cells were calculated based on the number of cells at a given time point that surpassed an arbitrary mAG1-geminin^1-110^ intensity threshold following palbociclib washout. At least 90 cells were analyzed for each condition at each time point. **(E)** The timing from G_1_ release (start of imaging) until mitotic entry for the first 40 cells that reach mitosis for each condition in the experiment described in (**D-F**). p-values were calculated by one-way ANOVA followed by Tukey’s multiple comparisons test. **** : p < 0.0001; ns : p = 0.9992. (**F**-**G**) Enlarged and size-constrained RPE1 cells expressing the FUCCI cell cycle markers (mCherry-Cdt1^30-120^ and mAG1-geminin^1-110^) were transfected with control or p53-directed siR-NAs as indicated in (**B**) and were imaged for 48 hours after drug washout. (**F**) Quantification of mitotic duration and mitotic failure rates in enlarged and size-constrained RPE1 cells that reached mitosis with and without p53 knockdown. Mitotic duration was quantified as the time from NEB to flattening or division. At least 25 cells were quantified per condition. p-values are given for mitotic duration and were calculated by two-way ANOVA followed by Tukey’s multiple comparisons test. * : p = 0.03; ns : p ≥ 0.05. Error bars = mean ± SD. (**G**) Quantification of mitotic slippage and nuclear fragmentation rates indicated in (**F**) as a fraction of total cells that entered mitosis for each condition. (**H**) Representative images of cells 48 hours following G_1_ arrest release. Magenta fluorescence indicates mCherry-Cdt1^30-120^ fluorescence (G_1_ marker), whereas green indicates mAG1-geminin^1-110^ expression (S/G_2_/M marker). Scale bar = 100 μm. (**I**) Quantification of (**H**). S/G_2_/M cells were defined based on FUCCI markers. 3-4 images were analyzed per condition, and each image contained 80-200 cells. Error bars = mean ± SD.

Increased p21 expression is a frequent consequence of active p53 signaling [42, 43]. Though we were not able to detect p53 in size-constrained or enlarged RPE1 cells by western blot or by mass spectrometry, we and others [44] found that depletion of p53 completely abrogates p21 levels in large cells (**Figure 3B-3C**). Of note, we found that p21 expression in enlarged RPE1 cells does not require active ATM or ATR signaling (**Figure S3D**-**S3G**) demonstrating that p53 activation in enlarged G_1_ cells is not initially triggered by the canonical DNA damage response pathway [45].

Because p21 is an inhibitor of the G_1_/S transition [29, 46], we examined whether p53 depletion affects cell cycle entry in oversized cells. Indeed, p53 depletion eliminates the fraction of enlarged cells that fail to enter the cell cycle, hastens the G_1_/S transition (**Figure 1F, Figure 3D**), and eliminates the cell cycle delays we observed in enlarged cells that reach mitosis (**Figure 1G, Figure 3E**). In agreement with the notion that p21 is the critical p53 target that mediates persistent G_1_ arrest in enlarged cells, others have shown that p21 depletion promotes cell cycle entry in enlarged cells [44]. Moreover, we found that depleting p53 in large cells increases the frequency of mitotic failure relative to control-transfected large cells but has no effect on cell cycle progression in size-constrained cells (**Figure 3F-3G, Figure S3H**). This result indicates that p53 protects against catastrophic cell division failure in enlarged cells but is dispensable for at least one cell cycle in size-constrained cells. Together, these data indicate that enlarged cells are primed to progress through the cell cycle but are restrained by active p53 signaling.

Following one round of cell division, large cells almost all arrest in G_1_, likely reflecting irreparable damage accrued during the previous cell cycle. In contrast, siP53-transfected large cells continue to cycle. This occurs even in cells that have undergone mitotic catastrophes like nuclear fragmentation or mitotic slippage (**Figure 3H-3I**). Thus, in summary, p53 limits cell cycle entry and mitotic failure in enlarged RPE1 cells, but it is not sufficient to block cell cycle progression in all cells. p53 signaling also prevents subsequent cell cycle re-entry following an initial cell cycle failure, thereby limiting the propagation of unstable genomes. This is consistent with data from others showing that removal of p53 promotes long-term proliferation following Cdk4/6 inhibitor withdrawal [11, 47, 48].

### Cell cycle entry in enlarged MCF7 cells is blocked by p21

To understand whether the phenotypes we observed in enlarged RPE1s are shared in other cell types, we analyzed cell size and cell cycle entry dynamics in NALM6 and MCF7 cells following release from prolonged palbociclib-mediated G_1_ arrests. Both of these cell lines are Rb+ positive and susceptible to palbociclib-mediated G_1_ cell cycle arrests [49]. NALM6 cells fail to accumulate significant biomass during a palbociclib-mediated arrest: over 6 days, they increase in volume less than two-fold (**Figure S4A-S4B**), which is modest compared to what we observed in RPE1 cells. The relatively unaffected cell volume in NALM6 cells correlated with normal proliferation upon release (**Figure S4C**-**S4D**), consistent with the notion that proliferation defects upon release are due to increased cell size. Importantly, these data suggest that cells that limit biomass accumulation during a prolonged G_1_ arrest are resistant to senescence induced by Cdk4/6 inhibition.

In contrast to NALM6 cells, MCF7 cells significantly increase in cell size upon prolonged palbociclib treatment. Using the scheme shown in **Figure 1A**, we generated size-constrained MCF7 cells by co-treating cells with Torin1 (**Figure 4A**). Importantly, plating MCF7 cells at high confluence does not contact inhibit growth [50] and therefore cannot be used as a strategy for constraining size in these cells. Consistent with our results from RPE1 cells, we found that enlarged MCF7 cells proliferated less well in long-term growth assays relative to size-constrained cells (**Figure 4B-4C**). Still, unlike RPE1 cells, MCF7 cells that have grown large recover very little cyclin A2 expression following G_1_ release, suggesting that a majority of cells fail to re-enter the cell cycle (**Figure 4D**). EdU incorporation measurements confirmed that nearly three times as many size-constrained cells entered S-phase relative to enlarged cells following release (**Figure 4E, Figure S4E**-**S4F**)—a behavior that we previously observed in IMR90 fibroblasts [7]. Taken together, our findings in both RPE1 and MCF7 cells confirm that excessive G_1_ cell size causes long-term cell cycle failure.

**Figure 4:**
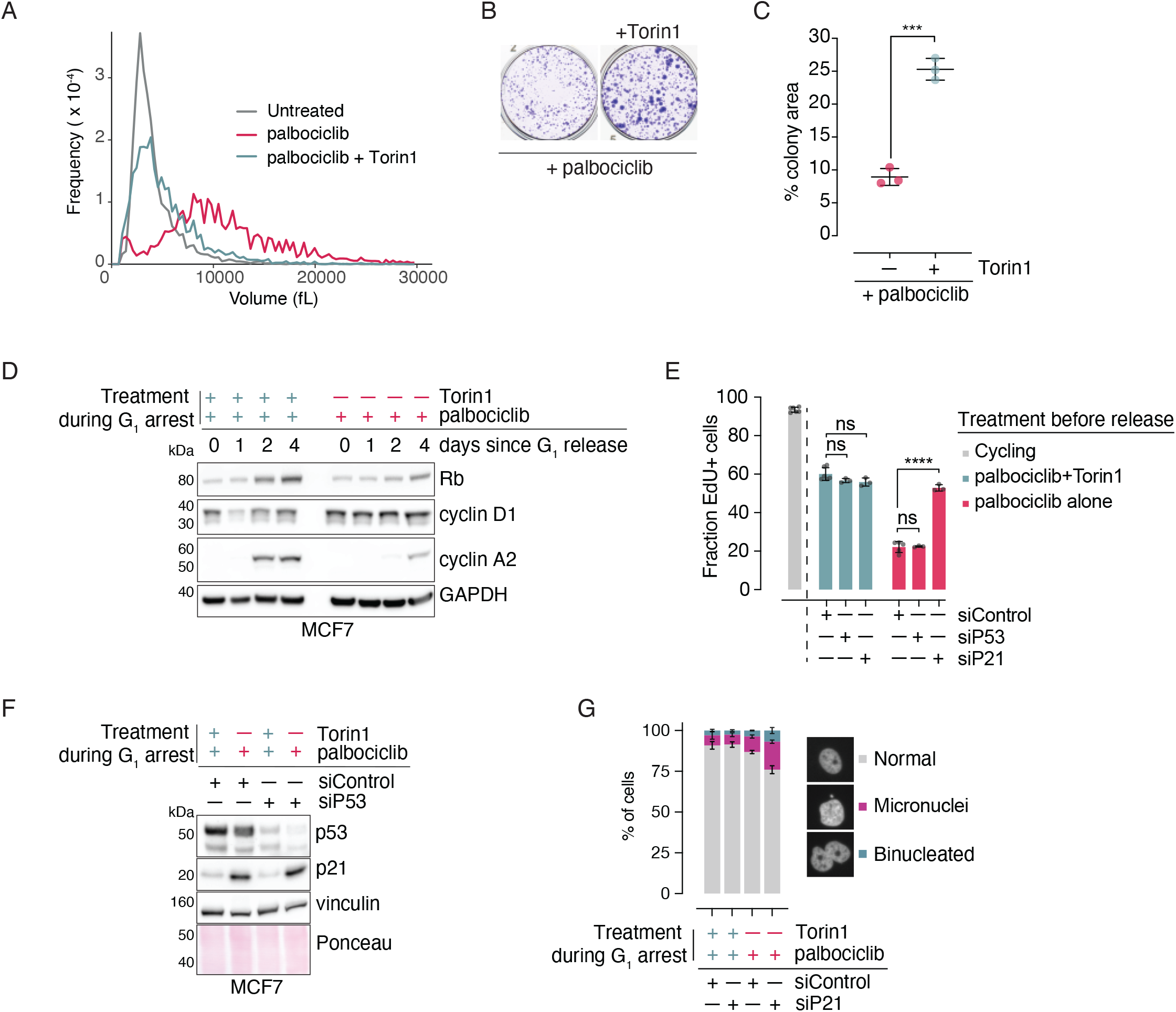
Cell cycle entry in enlarged MCF7 cells is blocked by p21. **(A)** Cell size measurements for cycling MCF7 cells and cells treated with palbociclib and/or Torin1 for 6 days. **(B)** Long-term colony formation assay for enlarged and size-constrained MCF7 released from G_1_ arrest. Cells were treated as in (**A**) with an additional day for recovery from Torin1-treatment (in the presence of palbociclib) and were then seeded at ∼250 cells/cm^2^ in the absence of drugs for 10 days. Cells were then fixed and stained with crystal violet to visualize colonies. **(C)** Quantification of (**B**). p-value was calculated by two-tailed, unpaired t-test. n=3. *** : p = 0.0002. Error bars = mean ± SD. **(D)** Western blots of whole cell lysates from a release time course following a 6-day G_1_ arrest in size-constrained (Torin1-treated) and enlarged MCF7 cells. After 6 days, cells were washed into palbociclib-only media for 24 hours before releasing into fresh media as indicated in **Figure 1A**. Cell lysates collected at the indicated time points and probed with the indicated antibodies. GAPDH was used as a loading control. **(E)** Size-constrained and enlarged MCF7 cells were treated with the indicated siRNA for 24 hours in the continuous presence of palbociclib (as shown in **Figure 3B**) before releasing into EdU-containing drug-free media for three days. Cells were then collected, and EdU was derivatized with AlexaFluor488. EdU incorporation was then measured by flow cytometry. At least three replicates were analyzed for each condition. ns : p ≥ 0.05, **** : p < 0.0001. p-values were calculated by two-way ANOVA followed by Tukey’s multiple comparison test. Error bars = mean ± SD. **(F)** Western blot depicting the results of p53 knockdown on p53 and p21 levels in size-constrained (Torin1-treated) and enlarged G_1_-arrested MCF7 cells. Vinculin and Ponceau staining were used as loading controls. **(G)** Size-constrained (Torin1-treated) and enlarged MCF7 cells were treated as in (**E**) using a p21-directed siRNA for 24 hours in the presence of palbociclib prior to drug washout. Two days after release, cells were fixed, nuclei were stained with Hoechst 33342, and nuclear defects were imaged by high-content fluorescence microscopy. The fraction of micronuclei and binucleated cells observed in each condition were calculated. Four replicates were measured for each condition, with approximately 300 cells per replicate. For simplicity, cells that were binucleated and had micronuclei were only categorized as binucleated. Error bars = mean ± SD.

Like enlarged RPE1 cells, enlarged MCF7 cells have elevated p21 levels. Still, unlike in RPE1 cells, this is not affected by siRNA-mediated p53 depletion (**Figure 4F**). Consistent with this result, p21 knockdown rescued S-phase entry in enlarged MCF7 cells (**Figure 4E, Figure S4E**) whereas p53 knockdown had no effect on cell cycle progression (**Figure 4F, Figure S4F**). Thus, p21 is required for cell cycle arrest in enlarged MCF7 cells. p21 knockdown also significantly increased the frequency of micronuclei and binucleated cells (**Figure 4G, Figure S4G**) in enlarged MCF7 cells two days after G_1_ arrest release. Because these phenotypes arise from failed mitoses, these data and our findings in RPE1 cells demonstrate that cell cycle progression in enlarged cells causes genome instability, and p21 is essential for limiting these defects.

### Excess cell size dampens DNA damage-induced p53 signaling

Because p53-dependent DNA damage signaling is important for suppressing mitotic defects associated with failed DNA replication [51-53], we hypothesized that deficits in p53 signaling could potentially explain the high rate of mitotic failure we observed in enlarged cells that progress through S-phase (**Figure 1H, Figure 3D-E, Figure 4G**). To investigate p53 dynamics in enlarged G_1_ cells, we measured how they respond to exogenous sources of DNA damage. We found that enlarged G_1_ RPE1 cells accumulated less p53 over time relative to size-constrained cells upon doxorubicin treatment (**Figure 5A-5D**). A similar result was observed in MCF7 cells treated with doxorubicin (**Figure S5A**). Moreover, whereas size-constrained RPE1 cells display a time-dependent increase in p21 expression that correlates with p53 stabilization, p21 induction is blunted in enlarged cells (**Figure 5A, Figure 5C**). Importantly, size-constrained and enlarged RPE1 cells stabilize p53 to the same extent in the presence of nutlin-3a—an MDM2 inhibitor that stabilizes p53 by blocking its degradation [54]— suggesting that the inability of large cells to mount an adequate p53 response does not arise from defects in p53 synthesis or in p21 induction but may instead be DNA damage-specific (**Figure 5E**). Consistent with this, others have shown that *TP53* mRNA levels do not fluctuate in response to changes in cell size [12]. In summary, p53 stabilization and p21 induction as a consequence of DNA damage are compromised in enlarged G_1_ cells.

**Figure 5:**
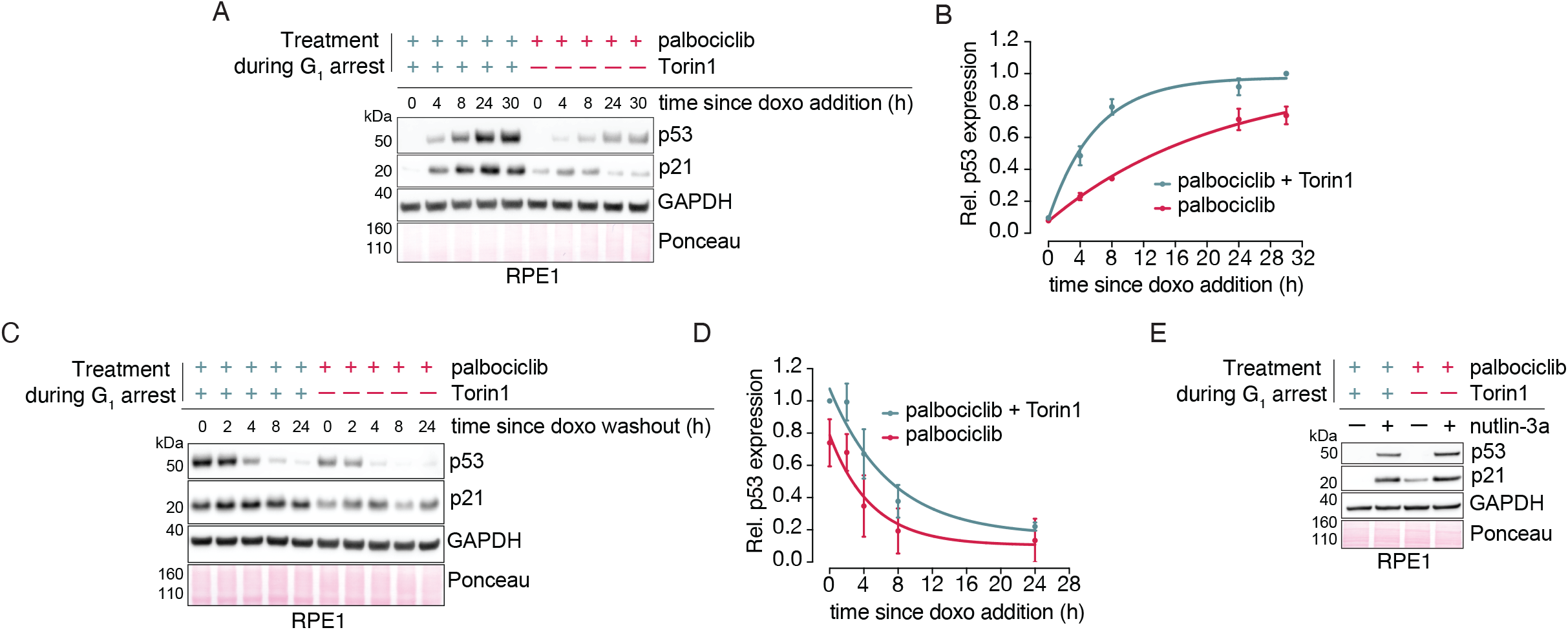
Excess cell size dampens DNA damage-induced p53 signaling. **(A)** RPE1 cells were treated as in **Figure 1A**, but the release step was omitted. After switching to recovery media (palbociclib alone) for one day, cells were treated with 1 μM doxorubicin in the continuous presence of palbociclib. Cells were collected at the indicated time points, and the indicated protein abundances were measured by western blot. GAPDH and Ponceau staining were used as loading controls. **(B)** Quantification of p53 levels shown in (**A**). Protein levels were normalized to GAPDH intensity and then the 30 hour size-constrained time point. Error bars = mean ± range for two experiments. **(C)** Enlarged and size-constrained RPE1 cells were treated with 500 nM doxorubicin in the presence of palbociclib for 16 hours before washing the doxorubicin out (maintaining the palbociclib) and taking samples for western blotting at the indicated time points. Protein abundances were measured by western blot using the indicated antibodies. GAPDH and Ponceau staining were used as loading controls. **(D)** Quantification of p53 levels from the experiment shown in (**C**). Protein levels were normalized to GAPDH intensity and then the 0 hr size-constrained time point. Error bars = mean ± range for two experiments. **(E)** Enlarged and size-constrained RPE1 cells were treated with 5 μM nutlin-3a in the presence of palbociclib for 24 hours. Cells were collected, and the indicated protein abundances were measured by western blot. GAPDH and Ponceau staining were used as loading controls.

### Enlarged G_1_ cells are prone to DNA damage

Because p53 plays an important role in driving cell cycle arrest and damage repair in response to DNA damage [13, 21, 28, 45], we asked whether excess cell size causes increased sensitivity to DNA damaging agents, which was recently reported in palbociclib-treated cells [11]. We found that the proliferation of enlarged cells released from G_1_ into a low dose of doxorubicin (**Figure 6A-6B**) or camptothecin (**Figure 6C-6D**) is stunted compared to size-constrained cells, indicating that DNA damage sensitivity following prolonged G_1_ arrest is a consequence of increased cell size and not the arrest itself. γH2AX staining demonstrated that enlarged G_1_ cells harbor higher levels of damage than size-constrained G_1_ cells following doxorubicin treatment (**Figure 6E-6F, Figure S6A**-**S6B**). This result was reproduced in enlarged MCF7 cells by western blotting (**Figure S6C**). Because H2AX is a histone and therefore subscales relative to cell size [12, 38] (**Figure 2B**), we normalized loading to histone content rather than total protein for this experiment. These results show that enlarged cells accumulate more damage in response to genotoxic stress compared to size-constrained cells. Because these experiments were carried out during a sustained G_1_ arrest, these data demonstrate that increased DNA damage sensitivity in enlarged cells is already established during G_1_. This likely contributes to the replication stress previously observed in palbociclib-treated cells [11]. Analysis of γH2AX foci in untreated cells revealed that enlarged G_1_-arrested cells have slightly elevated levels of basal DNA damage relative to size-constrained cells. Because p53 contributes to DNA damage repair, we asked whether the p53 defects we observed in enlarged cells could account for the increased level of damage we observed in enlarged cells. Removal of p53 in size-constrained cells modestly increases the number of cells with 1-2 γH2AX foci but was not sufficient to cause high levels of damage (e.g., cells with ≥3 foci). In contrast, p53 removal strongly exacerbated the level of severe endogenous DNA damage observed in enlarged cells (**Figure 6G**). Together, these data demonstrate that spontaneous DNA damage arises more frequently in enlarged G_1_ cells, and p53-dependent signaling— although inefficient— is required to repair it. Similarly, p53 knockdown is sufficient to increase the level of damage observed in both size-constrained and enlarged cells treated with doxorubicin (**Figure 6H, Figure S6D**), indicating that p53-dependent DNA damage repair reduces DNA damage levels during exposure to genotoxic agents. Importantly, enlarged cells wherein p53 is knocked down still harbor higher levels of damage than the corresponding size-constrained cells (**Figure 6G-6H, Figure S6D**). This indicates that—although p53 signaling defects may contribute to the high levels of DNA damage observed in enlarged cells— other factors also contribute to the high propensity for DNA damage accumulation observed in enlarged cells.

**Figure 6:**
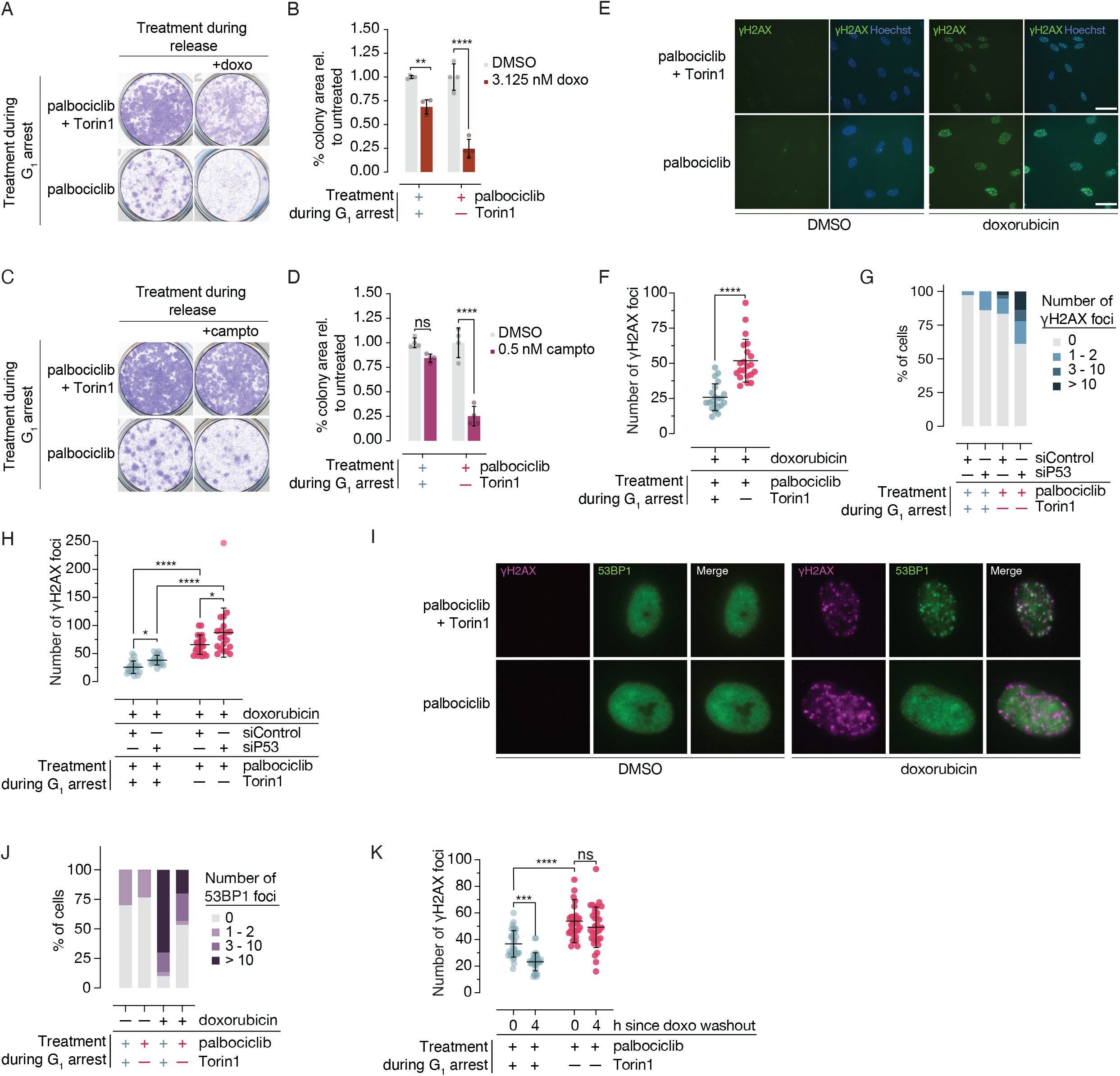
Enlarged G_1_ cells are prone to DNA damage. **(A)** RPE1 cells were treated as in **Figure 1A** and were re-seeded at ∼250 cells/cm^2^ in DMSO-containing media or 3.125 nM doxorubicin. Cells were then fixed and stained with crystal violet to visualize colonies after 10 days. **(B)** Quantification of (**A**). n = 4. p-values were calculated by two-way ANOVA followed by Tukey’s multiple comparison test. Samples were normalized to the DMSO treatment condition for each cell size condition. ** : p = 0.002 ; **** : p < 0.0001. Error bars = mean ± SD. **(C)** RPE1 cells were treated as in **Figure 1A** and were re-seeded at ∼250 cells/cm^2^ in DMSO-containing media or 0.5 nM camptothecin. Cells were then fixed and stained with crystal violet to visualize colonies after 10 days. **(D)** Quantification of (**C**). n = 4. p-values were calculated by two-way ANOVA followed by Tukey’s multiple comparison test. Samples were normalized to the DMSO treatment condition for each cell size condition ns : p = 0.2; **** : p < 0.0001. Error bars = mean ± SD. **(E)** Representative immunofluorescence images of γH2AX foci in size-constrained (palbociclib + Torin1) and enlarged (palbociclib) G_1_ RPE1 cells treated with DMSO or 1 μM doxorubicin for 24 hours. Hoechst 33342 was used as a nuclear counterstain. Scale bar = 70 μm. **(F)** Quantification of the doxorubicin conditions in the experiment shown in (**E**). Foci for at least 20 cells were counted for each condition. p-value was calculated by an unpaired two-tailed t-test. **** : p < 0.0001. Error bars = mean ± SD. **(G)** DMSO-treated enlarged and size-constrained cells (+/- p53 knockdown) were fixed and subjected to γH2AX immunofluorescence. γH2AX foci were counted and binned as indicated. At least 30 cells were analyzed for each condition. **(H)** Enlarged and size-constrained cells (+/- p53 knockdown) were treated with 1 μM doxorubicin for 24 hours in the continuous presence of palbociclib were immunostained for γH2AX. Foci were counted for each condition (20 cells each). p-values were calculated by two-way ANOVA followed by Tukey’s multiple comparisons test. * : p = 0.01 ; **** : p < 0.0001. (**I**-**J**) (**I**) Representative images of enlarged and size-constrained RPE1 cells that were treated as in (**E**-**F**), fixed, and subjected to γH2AX and 53BP1 immunofluorescence. (**J**) Quantification of the experiment described in (**I**). 53BP1 foci were counted and binned as indicated. At least 30 cells were analyzed for each condition. (**K**) Quantification of γH2AX staining in enlarged (palbociclib) and size-constrained (palbo-ciclib+Torin1) G_1_ arrested RPE1 cells that were treated with 1 μM doxorubicin for 16 hours (time = 0) and 4 hours after doxorubicin washout (remaining in palbociclib). Hoechst 33342 was used as a nuclear counter stain. At least 30 cells were analyzed for each condition. p-values were calculated by two-way ANOVA followed by Tukey’s multiple comparisons test. *** : p = 0.0002; **** : p < 0.0001 ; ns : p = 0.5.

The elevated γH2AX levels we observed in enlarged cells upon doxorubicin treatment are ATM-dependent (**Figure S6E**), indicating that doxorubicin causes double-stranded breaks (DSBs) [55]. We therefore investigated whether the DSB repair pathway was robustly maintained in enlarged cells. During G_1_, DSBs are repaired via non-homologous end-joining (NHEJ), which involves the formation of discrete 53BP1 foci at damage sites [56]. These foci then act as adaptors for downstream signaling [57]. To probe the DSB repair pathway in enlarged G_1_ cells, we measured 53BP1 foci formation upon doxorubicin treatment. We found that a significant portion of doxorubicin-treated enlarged cells fail to form 53BP1 foci, whereas size-constrained cells do so proficiently (**Figure 6I-6J**). Thus, an upstream component of the DSB repair pathway is impaired in enlarged cells. Moreover, because 53BP1 regulates the accumulation of p53 in response to DSBs [56], enlarged cells’ failure to stabilize p53 upon damage may be explained by deficits in 53BP1 foci formation.

To investigate whether the repair defects we observed translate to inefficient damage resolution, we treated enlarged and size-constrained cells with doxorubicin for 16 hours followed by a 4-hour washout and measured γH2AX foci. We found that size-constrained cells significantly reduce damage following doxorubicin removal, whereas enlarged cells fail to do so (**Figure 6K, Figure S6F**). Thus, impaired damage repair pathways in enlarged cells correlate with high levels of DNA damage and less efficient damage resolution. Together, these data show that blunted DNA damage signaling and an increased propensity for DNA breakage are established prior to S-phase in enlarged cells. We propose that this renders large cells hypersensitive to exogenous genotoxic insults and endogenous sources of damage like reactive oxygen species, transcription, and DNA replication.

## Discussion

In this study, we used reversible G_1_ cell cycle arrests coupled with different strategies for constraining cell size to study the effects of excess cell size on mammalian cell cycle progression. Based on this experimental system, we describe how unchecked growth in G_1_ causes cell cycle failure in non-transformed and transformed cell lines. Increased cell size drives the expression of p21, a Cdk1/2 inhibitor that signals G_1_ cell cycle arrest. In enlarged RPE1 cells, this is not sufficient to block cell cycle progression in all cells, whereas this is sufficient to block S-phase entry in the majority of enlarged MCF7 cells. In both cases, cell cycle progression through S-phase in enlarged cells causes increased genomic instability resulting from mitotic failures. Both cell lines fail to induce p53 robustly following DNA damage, demonstrating defects in a major pathway that drives p21 expression. We further show that excess size renders cells prone to accumulating DNA damage and interferes with efficient DNA damage repair, making enlarged cells highly sensitive to DNA damage.

We find that enlarged G_1_ RPE1 cells display heterogeneous cell cycle entry dynamics: whereas some cells are able to enter the cell cycle, others undergo prolonged G_1_ arrests. The population of cells wherein cell cycle entry is delayed is eliminated upon removal of p53, suggesting that differences in p53 signaling may account for the heterogeneity we observe in RPE1 cells. Based on the p53 signaling defects we observe in a bulk measurement of enlarged cells, these data fit a model in which some cells maintain robust p53 signaling, resulting in G_1_ cell cycle arrest, whereas others display weakened p53 signaling and enter the cell cycle. This is consistent with observations from others demonstrating that p53-dependent p21 expression is heterogeneous in RPE1 cells [58]. This model may also explain the discrepancy between the cell cycle entry behaviors we observed in RPE1 and MCF7 cells. Whereas most RPE1 cells enter the cell cycle, the majority of enlarged MCF7 cells fail to progress to S-phase. Though both cell lines harbor p53 signaling defects and elevated p21 levels, p21 expression in MCF7 cells is not affected by p53 depletion. Thus, MCF7 cells may be able to upregulate p21 more robustly— even in the absence of adequate p53 signaling—thereby resulting in a greater degree of G_1_ cell cycle arrest. Together, our findings from both cell lines indicate that p21 expression protects against inappropriate cell cycle entry and subsequent mitotic failures in enlarged cells.

Our proteomic analysis of G_1_ arrested large cells revealed that large cells have high protein levels of E2F transcriptional targets involved in DNA replication relative to size-constrained cells. Thus, DNA replication machinery is not limiting for proliferation in excessively large cells. Because p21—which should repress E2F gene expression through the inhibition of cyclin:Cdk complexes [13]— is elevated in enlarged cells, this suggests that opposing mechanisms drive E2F protein expression. This may be related to the recent observation that Rb concentrations are reduced as cells grow larger [19, 20]. In cells that have undergone unchecked G_1_ growth, Rb dilution may allow the untimely de-repression of E2F target genes. Such a mechanism would be independent of p21 levels, which inhibits E2F indirectly through Rb. Alternatively, high p21 levels may be overcome by the high levels of Cdk4 and cyclin D1 we observed in enlarged cells (**Figure S2B**), which others have shown is sufficient to accelerate the E2F transcriptional program [27]. Lastly, because Torin1-treatment and contact inhibition both limit mTOR signaling [59], growth restriction may repress E2F target expression in size-constrained cells. Indeed, others have found that a combination of Cdk4/6 inhibition and mTORC1/2 inhibition represses E2F-mediated transcription more so than Cdk4/6 inhibition alone [60], and recent proteomic measurements of rapamycin-treated cells revealed the suppression of DNA replication-related proteins [61].

We show that excess G_1_ cell size impairs DNA damage-induced activation of the p53 pathway, which contributes to inefficient DNA damage repair and high levels of damage in enlarged cells. We also found that enlarged cells fail to form 53BP1 foci in response to doxorubicin, whereas size-constrained cells do so proficiently. These results are consistent with previous observations that aged G_1_ cells—which are larger than healthy cells [62]—also fail to recruit 53BP1 to DSB sites [63]. This may be due to altered epigenetic conditions in enlarged cells given that 53BP1 foci formation requires specific histone modifications [57]. Alternatively, nuclear dilution could impede 53BP1 foci formation [64]. Because 53BP1 has been reported to enhance p53 signaling [56, 64-66], the failure to form 53BP1 foci at damage sites may contribute to the blunted p53 response we observe in enlarged cells.

When cells grow beyond their physiological size, macromolecule production becomes limited by DNA template availability [7]. Our observation that enlarged cells fail to robustly signal through DNA damage pathways suggests that cellular processes that generate signals from DNA itself (like DNA damage recognition and repair) may also become limited when the concentration of DNA decreases relative to cell volume. This has also been observed in the context of the spindle assembly checkpoint (SAC), where the kinetochore to cytoplasm ratio determines the strength of SAC signaling [67, 68]. Thus, the effect of cell size on other DNA-dependent processes should be studied further and may provide additional insight into why enlarged cells lose proliferative potential.

Others have hypothesized that Cdk4/6 inhibition induces replication stress due to downregulation of replisome components and origin licensing defects [11]. In agreement with these observations, our data demonstrate that replisome components are depleted over time during G_1_ arrest. Still, size-constrained cells contain even lower levels of replisome components than enlarged cells but have significantly lower rates of cell cycle failure upon G_1_ arrest release. Our data therefore do not support a model wherein replication machinery becomes limiting for DNA replication in enlarged cells. This finding reopens the question of why enlarged cells encounter replication stress during S-phase. Our data indicate that the DNA in enlarged G_1_ cells is damage-prone, due at least in part to defects in DNA damage repair signaling. Thus, the DNA damage sensitivity that we and others [11, 12] have observed in enlarged cells likely arises independent of DNA replication. We propose that DNA replication—an inherently damage-prone process [69]— is a general genotoxic stress that exposes the fragility of enlarged cells’ DNA in a manner similar to doxorubicin treatment. These defects may lead to persistent replication-induced breaks that would normally be mitigated, leading to high levels of irreparable DNA damage in cells entering G_2_. Defects in DNA damage-dependent p53 signaling may allow enlarged cells to enter mitosis in the presence of unresolved replication-acquired damage, culminating in failed mitoses and permanent cell cycle exit.

Lastly, our results present important clinical implications for the mechanism by which Cdk4/6 inhibitors induce permanent cell cycle withdrawal in cancer cells. Palbociclib and other Cdk4/6 inhibitors are frequently deployed as a treatment for HR+ and HER2– breast cancers [70]. Though others have shown that prolonged Cdk4/6 inhibition causes cell cycle failure [11, 44, 47, 50, 71], our data demonstrate that this is a consequence of increased cell size and is not strictly due to Cdk4/6 inhibition. This distinction suggests that Cdk4/6 inhibition may be a more useful therapeutic strategy for tumors containing cells that are susceptible to unchecked biomass accumulation. Consistent with this notion, others have shown that oncogenic mutations that amplify cell growth sensitize cells to palbociclib treatment [50], and hyperactivation of mTOR (which also amplifies cell growth) sensitizes ER+ breast cancer cells to Cdk4/6 inhibition in terms of permanent cell cycle withdrawal [72]. Conversely, our findings suggest that Cdk4/6 inhibition may not be a useful strategy for treating tumors wherein biomass accumulation is limited physically or through changes in signaling, as we observed in NALM6 cells. Lastly, because the loss of p53 in enlarged cells causes chromosome segregation defects but also allows continued cell cycle progression, our results suggest that Cdk4/6 inhibition may worsen chromosome instability in p53-null tumors wherein cells are able to grow large. Indeed, others have found that removal of p53 supports long-term proliferation following Cdk4/6 inhibitor withdrawal [47]. Thus, this work identifies potential limitations for using Cdk4/6 inhibitors in a therapeutic context and suggests caution in drawing clinical conclusions from *in vitro* experiments using these drugs.

## STAR Methods

### Key Resources Table

| REAGENT or RESOURCE | SOURCE | IDENTIFIER |
| --- | --- | --- |
| **Antibodies** |  |  |
| Rb | Cell Signaling Technology | #9309 |
| cyclin A2 | Santa Cruz | sc-751 |
| GAPDH | Abcam | ab8245 |
| vinculin | Santa Cruz | sc-73614 |
| p21 | Cell Signaling Technology | #2947 |
| MCM2 | Cell Signaling Technology | #3619 |
| cyclin D1 | Abcam | ab18521 |
| p16 | Cell Signaling Technology | #80772 |
| phospho-histone H2AX (rabbit) | Cell Signaling Technology | #2566 |
| phospho-histone H2AX (mouse) | Millipore | 05-636 |
| histone H3 | Abcam | Ab18521 |
| p53 | Santa Cruz | Sc-126 |
| Anti-mouse IgG HRP | BioRad | 170-6516 |
| Anti-rabbit IgG HRP | BioRad | 170-6515 |
| Anti-rabbit AlexaFluor488 | Thermo Fisher |  |
| Anti-mouse AlexaFluor568 | Thermo Fisher |  |
| **Chemicals, peptides, and recombinant proteins** |  |  |
| palbociclib | Sigma | PZ0383 |
| Torin1 | Sigma | 475991 |
| doxorubicin | Sigma | D1515 |
| camptothecin | Sigma | 208925 |
| KU55933 (ATM inhibitor) | Sigma | 1109 |
| AZ20 (ATR inhibitor) | Selleckchem | S7050 |
| nutlin-3a | Sigma | SML0580 |
| Hoechst 33342 | Thermo Fisher | H3570 |
| FxCycle FarRed DNA Stain | Thermo Fisher | F10348 |
| Lipofectamine RNAiMax | Thermo Fisher | 13778100 |
| NuPAGE LDS sample buffer (4x) | Thermo Fisher | NP0007 |
| Pierce protease and phosphatase inhibitor mini tablets | Thermo Fisher | A32959 |
| Pierce RIPA lysis buffer | Thermo Fisher | 89900 |
| 20x MES SDS running buffer | Thermo Fisher | B000102 |
| 20x MOPS SDS running buffer | Thermo Fisher | B0002 |
| **Critical commercial assays** |  |  |
| Click-iT EdU AlexaFluor-488 Flow Cytometry Assay Kit | Thermo Fisher | C10420 |
| Subcellular Protein Fractionation Kit for Cultured Cells | Thermo Fisher | 78840 |
| TMTpro 16plex | Thermo Fisher | 44522 |
| **Deposited data** |  |  |
| TMT proteomics in enlarged/size-constrained cells | This paper | PRIDE: PXD034934 |
| **Experimental models: Cell lines** |  |  |
| hTERT RPE1 WT | ATCC | CRL-4000 |
| hTERT RPE1 FUCCI | Laboratory of Randy King | N/A |
| MCF7 | ATCC | HTB-22 |
| NALM6 | Laboratory of Mike Tyers | N/A |
| **Oligonucleotides** |  |  |
| ON-TARGETplus Human TP53 | Dharmacon | J-003329-14-0002 |
| ON-TARGETplus Human CDKN1A | Dharmacon | J-003471-09-0002 |
| siGENOME non-targeting siRNA #5 | Dharmacon | D-001210-05-05 |
| **Software and algorithms** |  |  |
| FIJI | https://imagej.net/software/fiji/ | N/A |
| R | https://www.r-project.org/ | N/A |
| FlowJo | BD Biosciences | N/A |
| MaxQuant | Laboratory of Matthias Mann | N/A |
| ShinyGO 0.76 | http://bioinformatics.sdstate.edu/go/ | N/A |

#### Cell culture, growth conditions, and drug treatments

Cell lines used in this work (hTERT-RPE1 WT, hTERT-RPE1 FUCCI, MCF7, NALM6) were cultured in a humidified incubator at 37°C in the presence of 5% CO_2_. Wild-type hTERT-RPE1 and MCF7 cells were obtained directly from ATCC. hTERT-RPE1 FUCCI cells were a gift from Randall W. King with permission from the RIKEN Institute. NALM6 cells were a gift from Mike Tyers.

RPE1 cells were cultured in DMEM/F12 with GlutaMAX (Gibco) + 10% FBS and 1% penicillin/streptomycin. RPE1 cells that were allowed to grow large were seeded at low density (5,000 – 10,000 cells/cm^2^) and treated with 1 μM palbociclib for 6 days. Cells that were allowed to grow large were maintained at subconfluency for the duration of the experiment. RPE1 cells that were size-constrained using Torin1 treatment were seeded at 30,000 cells/cm^2^ prior to treating cells with 1 μM palbociclib + 500 nM Torin1. For contact inhibition experiments, RPE1 cells were seeded at 79,000 cells/cm^2^ for 48 hours before treating cells with 1 μM palbociclib. Note that this is ∼95% confluency after 24 hours. Cells were seeded at this density and allowed to grow for 48 hours because seeding at higher densities caused them to slough off the dish. After 6 days, cells were re-seeded at 30,000 cells/cm^2^ in the presence of 1 μM palbociclib for an additional 24 hours to recover and re-attach. For release experiments, cells were then washed 3x in media and released into media without drugs.

MCF7 cells were cultured in DMEM with GlutaMAX (Gibco) + 10% FBS and 1% penicillin/streptomycin. MCF7 cells were seeded at 30,000 cells/cm^2^ prior to treating cells with 2 μM palbociclib for 6 days. To constrain cell size, MCF7 cells were co-treated with 12 nM Torin1. Note that MCF7 cells are highly sensitive to mTOR inhibition, and higher doses caused significant cell death. For release experiments, cells were washed 3x in media and released into media without drugs.

NALM6 cells were cultured in RPMI-1640 with GlutaMAX (Gibco) + 10% FBS and 1% penicillin/streptomycin. NALM6 cells were maintained between 100,000 cells/mL and 1 × 10^6^ cells/mL. Note that NALM6 cells are sensitive to different FBS sources and doubling time (∼24 hours) should be confirmed for a given FBS source before use.

#### siRNA transfections

Cells were reverse-transfected using Lipofectamine RNAiMax (Invitrogen 13778100) according to manufacturer’s instructions with the following siRNAs at a final concentration of 25 nM. Cells were transfected for 24 hours for all experiments. Knockdowns were confirmed by western blotting. For G_1_ arrest experiments, transfections were carried out in the constant presence of palbociclib.

#### Cell size measurements

Cell size was measured on a Multisizer 4e (Beckman) using Isotone II (Beckman) as a diluent and a 100 μm aperture. For experiments in RPE1 cells, trypsinized cells were resuspended in DMEM/F12 and diluted in 10 mL Isotone II before measuring. MCF7 cells were trypsinized and diluted in Isotone II without re-suspending in media to minimize clumping. NALM6 cells were directly measured in RPMI-1640 diluted in Isotone II. Cell volume distributions were analyzed using a custom R script.

#### Crystal violet staining-based colony formation assays

For colony formation experiments, cells were seeded at ∼260 cells/cm^2^ in the presence of palbociclib (1 μM for RPE1 cells; 2 μM for MCF7 cells) for 24 hours to re-attach. Cells were then gently washed 3x in media and released into fresh media. For doxorubicin and camptothecin sensitization experiments in RPE1 cells, cells were released into 3.125 nM doxorubicin or 0.5 nM camptothecin, respectively. Cells were allowed to grow for 10-12 days before washing 1x with ice-cold PBS followed by incubation with ice-cold 100% methanol for 10 minutes on ice. The methanol was aspirated, and then cells were incubated with 0.5% crystal violet (w/v) in 25% methanol at room temperature for 10 minutes. Cells were washed with DI water until clear. Plates were dried at room temperature prior to imaging. Colony formation was quantified using the automated ColonyArea macro in ImageJ [73].

#### SDS-PAGE and western blotting

Cells were lysed in RIPA lysis buffer (Invitrogen 89900) supplemented with 1x protease/phosphatase inhibitor tablets (Pierce A32959) by periodic vortexing on ice. Lysates were clarified by centrifugation at 13,000rpm for 10 minutes at 4°C. Lysates were transferred to fresh tubes, and protein concentrations were measured using a BCA assay kit according to manufacturer’s instructions (Pierce 23225). Lysates were combined with 4x LDS sample buffer (Invitrogen NP0007) + 25 mM DTT to a final concentration of 1x and were boiled at 95°C for 10 minutes. Equal masses of protein (except for histone-normalized experiments, where lysate was derived from an equal number of cells) were loaded on Bolt 4-12% Bis-Tris gels (Invitrogen) and resolved in either 1x MES or MOPS SDS running buffer (Invitrogen B000102, B0002). Proteins were transferred to PVDF membranes at 250 mA at 4°C for 1 hr 15 min using a wet transfer apparatus. Membranes were blocked in 5% milk in TBS-T before incubating with primary antibodies in 5% milk in TBS-T at 4°C with agitation overnight. Membranes were washed 3x in TBS-T for ∼10 minutes each, incubated with secondary mouse or rabbit IgG HRP-conjugated antibodies in 5% milk in TBS-T for 45-60 minutes, washed again, and then developed using enhanced chemiluminescent substrate solutions as described by the manufacturer (Thermo 34095, Thermo 34578).

#### Chromatin fractionation

Isolation of chromatin bound MCM2 and PCNA was carried out using a subcellular fractionationation kit for cultured cells (Thermo 78840) according to the manufacturer’s protocol. Isolated fractions were diluted to 1x in LDS sample buffer (Invitrogen NP0007) + 25 mM DTT and were analyzed by SDS-PAGE and western blotting as described above.

#### Time lapse fluorescence imaging

G_1_ arrested cells were plated in the presence of palbociclib on an 8-well coverslip dish (Ibidi 80826) to be approximately 70% confluent after 24 hours. For siRNA transfection experiments, cells were transfected at the time of plating on coverslip dishes. For cell cycle release experiments, compounds (and siRNAs) were washed out and cells were imaged immediately following the addition of fresh media. Coverslips were inserted into a covered cage microscope incubator (Okolabs) with temperature and humidity control at 37°C and 5% CO_2_ and mounted on a motorized microscope stage (Prior ProScan HLD117NN). All images were collected on a Nikon Ti motorized inverted microscope equipped with a piezo z-drive (Prior), a 20x/0.75 NA S Fluor air objective lens, and the Perfect Focus system. mCherry fluorescence was excited with a Lumencor Spectra III light engine using a 555/28 excitation filter and a 641/75 emission filter (Semrock). mAG1 fluorescence was excited using a 475/28 excitation filter and a 515/30 emission filter (Semrock). Images were acquired with an Orca Fusion BT camera controlled with Nikon Elements image acquisition software. Four fields of view were collected per condition, and brightfield and/or fluorescence images were captured at 5-6 minute intervals.

#### Cell fixation and staining

##### Immunostaining and imaging

Cells were seeded onto glass coverslips and treated as indicated. Cells were then fixed with 4% formaldehyde solution for 10 min and permeabilized with 0.25% Triton X-100 in PBS for 10 min. Fixed cells were washed three times for 5 minutes with 0.05% Triton X-100 in PBS and blocked with 1% BSA in PBS for 30 min at room temperature. For staining of only γH2AX, cells were incubated with the primary antibody (CST #2577, 1:1000) in blocking solution for 1 hour at room temperature. Cells were later washed three times for 5 minutes with 0.05% Triton X-100 in PBS. For detection of the primary antibody, cells were incubated with an AlexaFluor488-conjugated secondary rabbit antibody (Thermo Fischer Scientific, 1:1000) in blocking solution for 1 hour at room temperature. For co-staining of γH2AX and 53BP1, cells were incubated with a γH2AX primary mouse antibody (Millipore 05-636, 1:500) and a 53BP1 primary rabbit antibody (Abcam ab21083, 1:400) in blocking solution for 1 hour at room temperature. For detection of the primary antibodies, cells were incubated with an AlexaFluor568-conjugated secondary mouse antibody (Thermo Fisher Scientific, 1:1000) and an AlexaFluor488-conjugated secondary rabbit antibody (1:400) in blocking solution for 1 hour at room temperature. Cells were then washed once with 0.05% Triton X-100 in PBS for 5 minutes and washed twice with 0.05% Triton X-100 + Hoechst 33342 (0.1 μg/mL) in PBS for 5 minutes. Coverslips were washed shortly with Milli-Q water and mounted onto glass slides using Vectashield mounting medium (Vector Laboratories).

Microscopy was performed on a Nikon Eclipse Ti-E inverted microscope using a 60x/NA 1.40 oil objective. Hoechst fluorescence was excited with a Lumencor SpectraX light engine using a 390/18 nm excitation filter and a 460/50 nm emission filter (AHF Analysentechnik). AlexaFluor488 fluorescence was excited using a 470/40 nm excitation filter and a 520/35 nm emission filter (AHF Analysentechnik). AlexaFluor568 fluorescence was excited using a 575/27 nm excitation filter and a 641/75 nm emission filter (AHF Analysentechnik). Images were acquired with a Hamamatsu ORCA Flash 4.0 camera controlled by the ImageJ μManager software [74]. Five to ten fields of view were collected per condition.

##### EdU staining and flow cytometry-based detection

MCF7 and NALM6 cells were washed out of palbociclib-containing media and into 10 μM EdU-alkyne for three days. Cells were then collected, fixed, and labeled with AlexaFluor-488-azide according to the manufacturer’s protocol (Thermo C10632). EdU incorporation was analyzed using a BD FACSCanto cell analyzer. Cells were co-stained for DNA using FxCycle FarRed (Thermo F10348) in the presence of RNase A to gate for single cells. Note that this experimental setup was not viable in RPE1 cells due to cell lysis during fixation and sample preparation.

##### Fixed cell imaging of nuclear abnormalities

For analysis of nuclear defects in MCF7 cells, enlarged and size-constrained MCF7 cells were seeded in a black-walled 96-well plate (Corning 3606) in the presence of palbociclib for 24 hours. Cells were then washed 3x with media and released from drug treatment for two days. Cells were then fixed (10% formalin and 0.1% Triton X-100 in PBS) and DNA was stained with Hoechst 33342 (1 μg/ml) for 45 minutes in the dark before imaging with an ImageXpress Micro high content microscope (Molecular Devices) equipped with a 20x objective and the DAPI filter. At least 15 images were collected and analyzed for each condition.

#### TMT mass spectrometry sample preparation

Samples were prepared essentially as described previously [75]: Cells were cultured as described in biological triplicate. Cell pellets were re-suspended in urea lysis buffer: 8M urea, 200 mm EPPS pH 8.0, Pierce protease inhibitor tablets (Thermo Fisher Scientific, A32963), and Pierce phosphatase inhibitor tablets (Thermo Fisher Scientific, A32957). Lysates were passed through a 21-gauge needle 20 times, and protein concentrations were measured by BCA assay (Thermo Fisher Scientific). One hundred micrograms of protein were reduced with 5 mm tris-2-carboxyethyl-phosphine (TCEP) at room temperature for 15 min, alkylated with 10 mm iodoacetamide at room temperature for 30 min in the dark, and were quenched with 15 mm DTT for 15 min at room temperature. Proteins were precipitated using a methanol/chloroform extraction. Pelleted proteins were resuspended in 100 µL 200 mm EPPS, pH 8.0. LysC (Wako 125-05061) was added at a 1:50 enzyme/protein ratio, and samples were incubated overnight at room temperature with agitation. Following overnight incubation, trypsin (Promega V5111) was added at a 1:100 enzyme/protein ratio, and samples were incubated for an additional 6 h at 37 °C. Tryptic digestion was halted by the addition of acetonitrile (ACN). Tandem mass tag (TMT) isobaric reagents (TMTpro 16plex Thermo Fisher Scientific 44522) were dissolved in anhydrous ACN to a final concentration of 20 mg/mL, of which a unique TMT label was added at a 2:1 label:peptide ratio. Peptides were incubated at room temperature for one hour with vortexing at the 30-minute interval. TMT labeling reactions were quenched by the addition of 10 µL of 5% hydroxylamine. Equal amounts of each sample were combined at a 1:1 ratio across all channels and dried by vacuum centrifugation. Samples were re-suspended in 1% formic acid and desalted using a 50 mg 1 cc SepPak C18 cartridge (Waters WAT054955) following manufacture’s instruction. Briefly, peptides were washed with 5% ACN and 0.1% formic acid, eluted with 50% ACN and 0.1% formic acid and dried. Subsequently, peptides were subjected to fractionation with basic pH reverse phase HPLC chromatography using a linear gradient (5-40% acetonitrile, 9mM ammonium bicarbonate) on XBridge peptide BEH C18 column (130 Å, 3.5 μm, 4.6 mm X 250 mm, Waters). Fractions were collected in 96 well format plate and consolidated on 12 fractions, dryed and re-suspended in 5% acetonitrile and 5% formic acid for LC-MS/MS processing.

#### TMT Mass Spectrometry Analysis

LC-MS/MS analysis was performed on an Orbitrap Fusion Lumos Tribrid mass spectrometer (Thermo Scientific) coupled to an Acquity UPLC M-class system (Waters). Peptides were loaded on a commercial trap column (Symmetry C18, 100Å, 5µm, 180 µm*20mm, Waters) and separated on a commercial column (HSS T3, 100Å, 1.8µm, 75 µm*250mm, Waters) using a 113 min gradient from 5% to 35% acetonitrile at a flow rate of 300 nL/min. The mass spectrometer was operated in data dependent acquisition (DDA) mode with 2s cycle time. MS1 data were collected in the Orbitrap (400-1400 m/z) at 60’000 resolution, 50 ms injection time and 4e5 AGC target. Ions with charge states between two and six were isolated in quadrupole (isolation window 0.5 m/z), fragmented (CID, NCE 35%) and MS2 scans were collected in the ion trap (Turbo, maximum injection time 120 ms, AGC 1.5e4); 60s of dynamic exclusion was used. MS3 quantification scans were performed with ten notches; ions were isolated in the quadrupole (2 m/z), fragmented (HCD, NCE 45%) and identified in the Orbitrap (50’000 resolution, maximum injection time 86ms and AGC 2e5).

#### Mass Spectrometry Data Analysis

Acquired spectra were searched using the MaxQuant software package version 2.1.0.0 embedded with the Andromeda search engine [76] against the human proteome reference dataset (http://www.uniprot.org/, downloaded on 06.04.2021) extended with reverse decoy sequences. The search parameters were set to include only full tryptic peptides (Trypsin/P), maximum two missed cleavage, carbamidomethyl and TMT16 as static peptide modification, oxidation (M) and acetylation (Protein N-term). Precursor and fragment ion tolerance was set respectively to 4.5ppm and 20ppm. False discovery rate of <1% was used at the PSM and protein level. Reporter intensities for proteins identified with at least 2 peptides (5884) were normalized, and missing values (1.7%) were imputed using random sampling from a normal distribution generated from 1% less intense values. ANOVA statistical tests were performed to compare protein profiles in all conditions. *P*-values were corrected using the Benjamini-Hochberg method [77]. Matrices with protein intensities and ANOVA statistical tests are reported in **Supplemental Data Table 1**.

#### Gene Enrichment Analysis

Gene enrichment analysis of the subset of proteins identified in the proteomics with a significant (adj. p-value < 0.05) increase in abundance with a log_2_(FC) ≥ 1 using both size-constraint methods was performed using the ShinyGO tool for biological processes [78].

## Supplemental Material

Supplemental Figures S1-S6

Supplemental Data Table 1 : TMT proteomics (full table)

Supplemental Data Table 2 : Decreasing proteins

Supplemental Data Table 3 : Increasing proteins

## Data availability

Mass spectrometry data have been deposited to the ProteomeXchange Consortium with the identifier PXD034934. All other raw data and reagents generated from this study are available from the corresponding author upon request.

## Acknowledgements

We are grateful to R. King and M. Tyers for sharing cell lines, M. Peter for sharing reagents and laboratory infrastructure, R. Meier and T. Schwarz from ScopeM for assistance with microscopy experiments, A. Dittmann and B. Roschitzki from Functional Genomics Center Zürich (FGCZ) for laboratory support and maintenance of the mass spectrometry facility. This work was supported by an ETH Postdoctoral Fellowship to S.M. (1-008383) and a SNSF Eccellenza grant to G.E.N. (PCEFP3_187003).

## Author Contributions

Conceptualization: S.M. and G.E.N.; Methodology: S.M., M.E.E., and F.U.; Investigation: S.M., M.E.E., and F.U.; Writing – Original Draft: S.M. and G.E.N.; Writing – Reviewing & Editing: S.M., M.E.E, F.U., and G.E.N.; Funding Acquisition: S.M. and G.E.N.; Supervision: G.E.N.

---

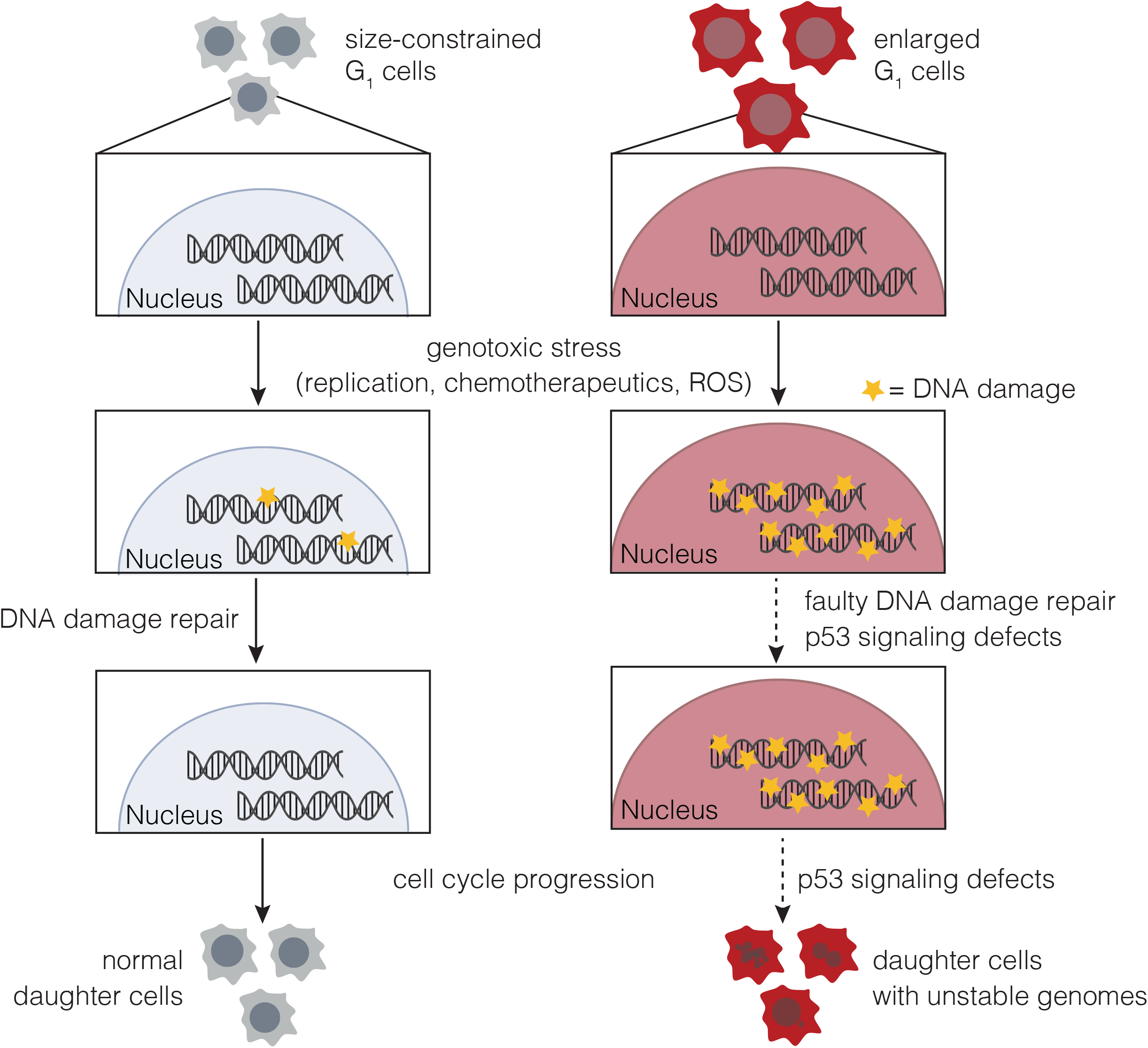

