## Supplemental Figure 1 for "Cell cycle progression defects and impaired DNA damage signaling drive enlarged cells into senescence"

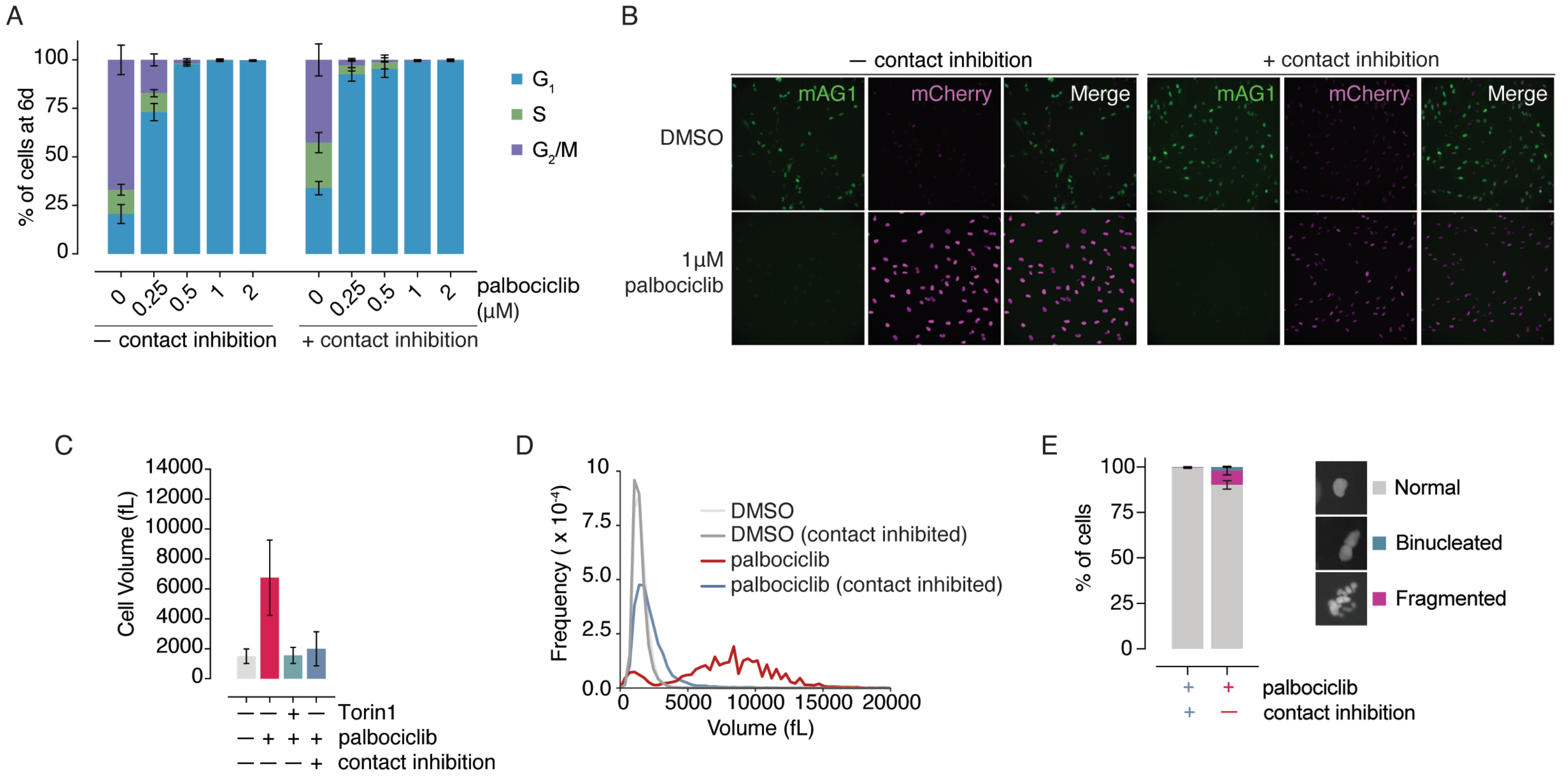


**Figure S1**. Additional data from Figure 1

(**A**) Cell cycle distributions for RPE1 FUCCI cells treated with various doses of palbociclib (+/- contact inhibition) for 6 days + 1 day of recovery as measured by FUCCI cell cycle reporters. Error bars = mean ± SD for three replicates. Four images were analyzed for each replicate with at total of at least 230 cells scored per replicate. 1 µM palbociclib was used for all of the RPE1 experiments in this study.
(**B**) Representative images of RPE1 FUCCI cells treated with DMSO or 1µM palbociclib for 6 days + 1 day of recovery as in (**A**).
(**C**) Cell size data from **Figure 1B** depicted as a bar plot. Error bars = mean ± SD.
(**D**) Coulter Counter based cell size measurements on Day 6 of the arrest for the experiment shown in **Figure 1H**.
(**E**) Fractions of enlarged and size-constrained RPE1 WT cells (contact inhibition) that were binucleated or fragmented six days after $G_1$ arrest release. At this time, cells were fixed, nuclei were stained with Hoechst 33342, and nuclear defects were imaged by high content fluorescence microscopy. The fraction of fragmented nuclei and binucleated cells observed in each condition was calculated. Three replicates were measured for each condition, with at least 450 cells scored per replicate. Error bars = mean ± SD.
