## Supplemental Figure 2 for "Cell cycle progression defects and impaired DNA damage signaling drive enlarged cells into senescence"

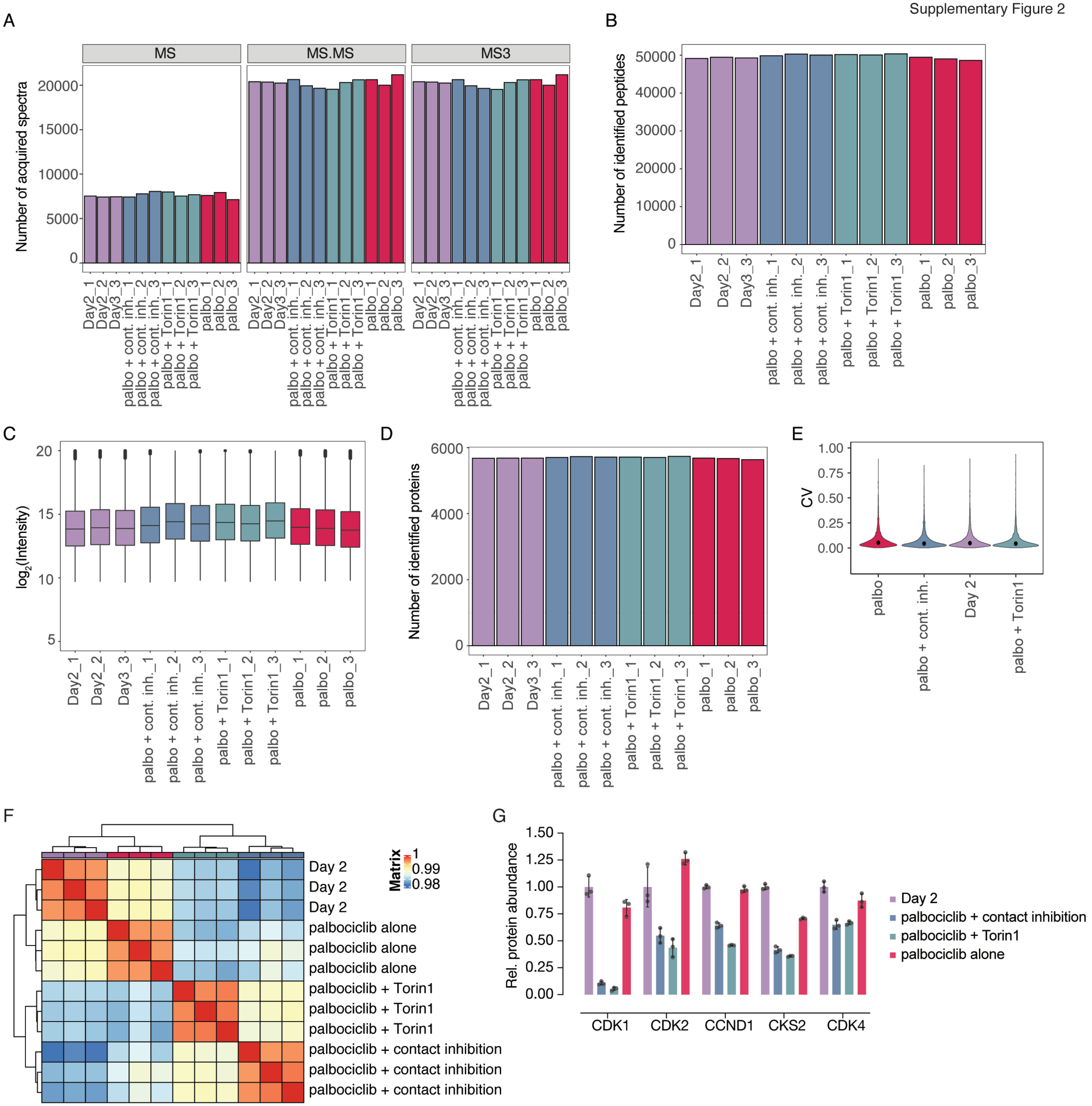


**Figure S2**. Additional data from Figure 2

(**A**) Number of spectra acquired in MS1, MS2, and MS3 for each reporter ion channel per acquired run for the $G_1$ cell size TMT proteomics experiment described in Figure 2.
(**B**) Number of peptides identified per reporter ion channel.
(**C**) Boxplot of peptide intensities per reporter ion channel.
(**D**) Number of proteins identified per reporter ion channel.
(**E**) Coefficient of variation for protein intensities in each condition.
(**F**) Unsupervised hierarchical clustering analysis of replicates.
(**G**) Protein abundances for various positive regulators of the $G_1$/S transition as measured by mass spectrometry. Protein abundances were normalized to the Day 2 samples. Error bars = mean ± SD.
