## Supplemental Figure 3 for "Cell cycle progression defects and impaired DNA damage signaling drive enlarged cells into senescence"

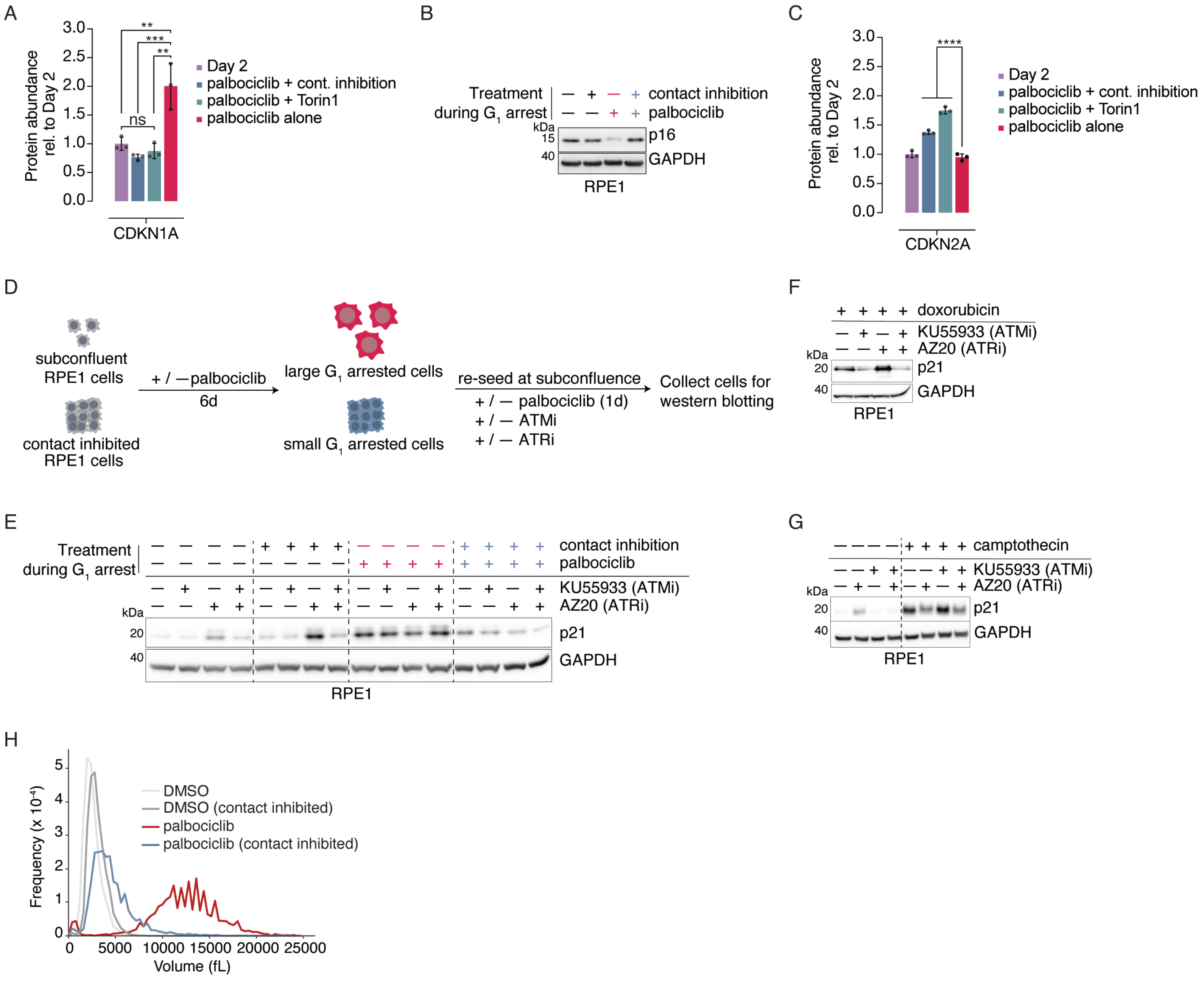


**Figure S3.** Additional data from Figure 3

(**A**) Quantification of p21 protein abundance based on MS measurements in Day 2, Day 6 size-constrained (palbociclib + either contact inhibition or Torin1-treated), and Day 6 enlarged $G_1$ cells (palbociclib alone). Protein abundances were normalized to Day 2 samples. p-values were calculated by one-way ANOVA followed by Tukey's multiple comparisons test. Error bars = mean ± SD. ** : $p < 0.01$; *** : $p = 0.0006$. ns : $p > 0.05$.
(**B**) Measurement of p16 levels in cycling, size-constrained (contact inhibited) and enlarged RPE1 cells. GAPDH was used as a loading control.
(**C**) Quantification of p16 protein abundance based on MS measurements. Proteins were normalized to Day 2 samples. p-values were calculated by one-way ANOVA followed by Tukey's multiple comparisons test. Error bars = mean ± SD. **** : $p < 0.0001$.
(**D**) Schematic for measuring the effect of ATM and ATR signaling on p21 levels in enlarged and size-constrained RPE1 cells. Small molecules KU55933 and AZ20 were used to block ATM and ATR activity, respectively.
(**E**) Experiment conducted with the schematic shown in (**D**). p21 levels were measured by western blotting, and GAPDH was used as a loading control.
(**F**) Validation that KU55933 can lower p21 levels as a function of ATM inhibition in RPE1 cells. Cycling RPE1 cells were treated with 1 µM doxorubicin +/- KU55933 and/or AZ20 24 hours before cells were collected and the indicated protein abundances were measured by western blotting. GAPDH was used as a loading control.
(**G**) Validation that AZ20 can lower p21 levels as a function of ATR inhibition in RPE1 cells. Cycling RPE1 cells were treated with or without 5 µM camptothecin and/or KU55933 and/or AZ20 for 24 hours, after which point cells were collected and the indicated protein abundances were measured by western blotting. GAPDH was used as a loading control.
(**H**) Coulter Counter based cell size measurements on Day 6 of the arrest (at the time of siRNA transfection) for the experiment shown in **Figure 3F-3G**.
