## Supplemental Figure 4 for "Cell cycle progression defects and impaired DNA damage signaling drive enlarged cells into senescence"

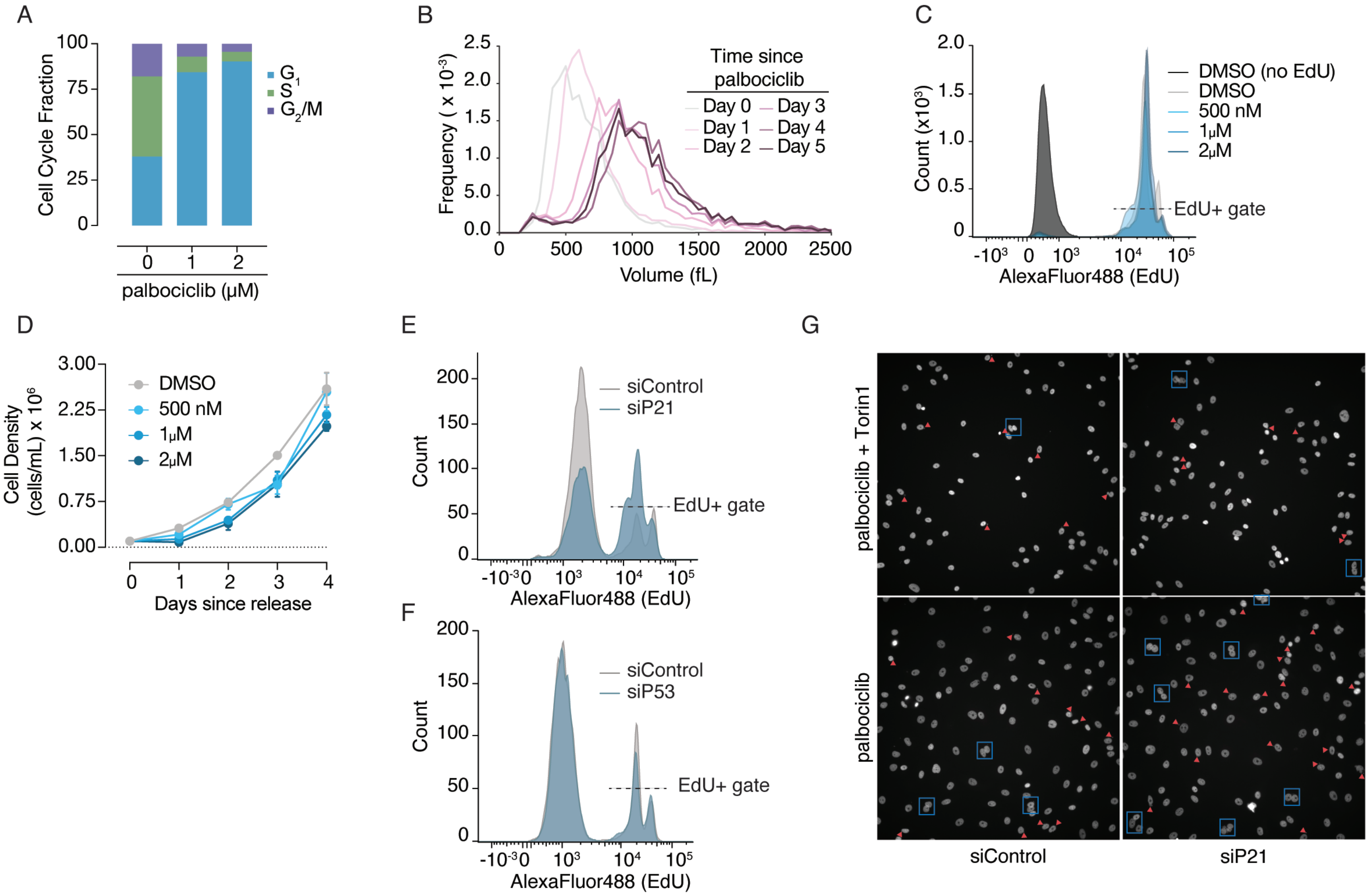


**Figure S4.** Additional data from Figure 4

(**A**) NALM6 cells were treated with the indicated doses of palbociclib for four days, after which cells were fixed and DNA was stained with FxCycle FarRed. Cell cycle profiles were obtained by measuring FxCycle FarRed DNA stain incorporation by flow cytometry.
(**B**) NALM6 cells were treated with 1 µM palbociclib for five days, and cell volume was measured each day using a Coulter Counter. Note that cells were maintained at 300,000 cells/mL for the duration of the experiment, though cell number only changed between Day 0 and Day 1. Day 0 indicates the time at which palbociclib was added.
(**C**) NALM6 cells were treated with the indicated concentrations of palbociclib for six days, and were then released into EdU-containing media in the absence of palbociclib for an additional three days. Cells were then fixed, and EdU was conjugated to AlexaFluor-488, followed by flow cytometry. The dashed line indicates EdU instensities that were considered EdU+ cells.
(**D**) NALM6 cells were treated with the indicated doses of palbociclib for six days as shown in (**C**). Cells were then spun down and re-plated in drug free media at a density of 100,000 cells/mL. Cells were then counted each day for four days following release. Error bars = mean ± SD.
(**E** and **F**) Representative EdU incorporation histograms for the released, enlarged (palbociclib alone) MCF7 cells following p21 (**E**) and p53 (**F**) knockdown in the experiment described in **Figure 4E**. The dashed lines indicate cells that were considered EdU+.
(**G**) Representative images from the experiment described in **Figure 4G**. Blue boxes show binucleated cells and red triangles point to micronucleated cells.
