## Supplemental Figure 5 for "Cell cycle progression defects and impaired DNA damage signaling drive enlarged cells into senescence"

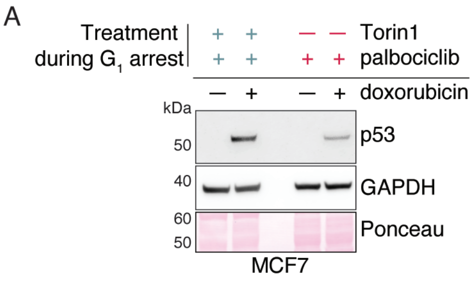


**Figure S5.** Additional data from Figure 5

(**A**) MCF7 cells were treated as in **Figure 1A** using Torin1 to constrain cell size, but the release step was omitted. After switching to recover media (palbociclib alone) for one day, cells were treated with 500 nM doxorubicin in the continuous presence of palbociclib. Cells were collected after 24 hours, and p53 levels were meausred by western blot. GAPDH and Ponceau staining were used as loading controls.
