## Supplemental Figure 6 for "Cell cycle progression defects and impaired DNA damage signaling drive enlarged cells into senescence"

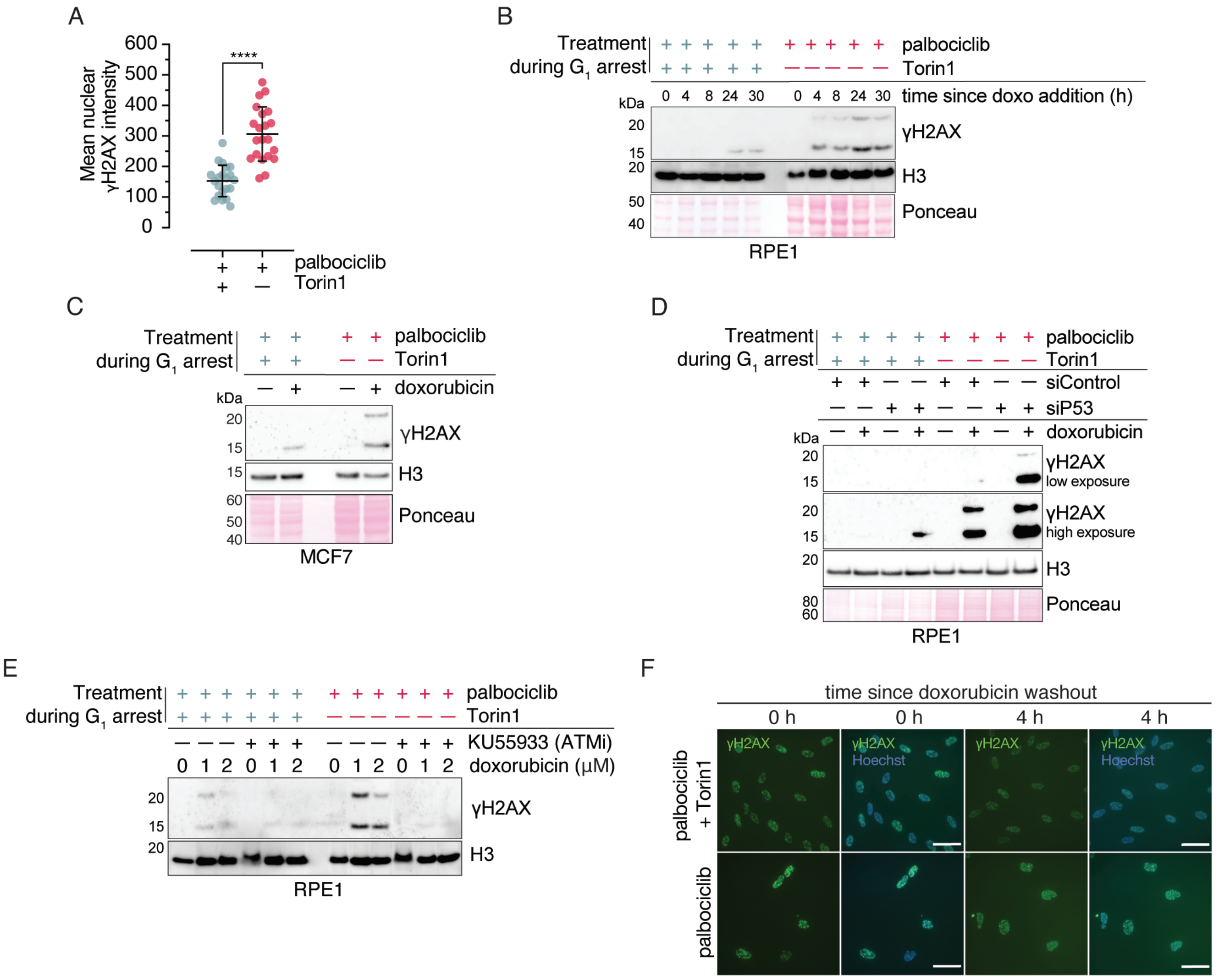


**Figure S6.** Additional data from Figure 6

(**A**) The experiment from **Figure 6E-6F** was alternately analyzed by measuring background-subtracted mean nuclear γH2AX intensity for the same cells shown in **Figure 6F**. p-value was calculated by an unpaired two-tailed t-test. **** : $p < 0.0001$. Error bars = mean ± SD.
(**B**) Measurement of γH2AX levels following doxorubicin treatment in RPE1 cells by western blot. Cells were treated as in **Figure 5A**, however lysates were normalized to cell number instead of protein amount. Histone H3 was used as a loading control to normalize to histone content, and Ponceau staining was used to illustrate differences in total protein abundance.
(**C**) Enlarged and size-constrained MCF7 cells were treated as in **Figure 6E-6F**, with the exception that 500 nM doxorubicin was used instead of 1 μM. Equal numbers of cells were lysed, and the indicated protein abundances were measured by western blotting. Histone H3 was used as a loading control to normalize to histone content, and Ponceau staining was used to illustrate differences in total protein abundance.
(**D**) Measurement of γH2AX levels by western blot for the experiment shown in **Figure 6H**. Histone H3 was used as a loading control to normalize to histone content, and Ponceau staining was used to illustrate differences in total protein abundance.
(**E**) Enlarged and size-constrained RPE1 cells were treated with the indicated doses of doxorubicin and KU55933 for 24 hours in the continuous presence of palbociclib. Equal numbers of cells were lysed, and the indicated protein abundances were measured by western blotting. Histone H3 was used as a loading control to normalize to histone content.
(**F**) Representative images from the experiment described in **Figure 6K**. Hoechst 33342 was used as a nuclear counterstain. Scale bar = 70 μm.
